# Identification, crystallization and epitope determination of public TCR shared and expanded in COVID-19 patients

**DOI:** 10.1101/2021.03.23.436573

**Authors:** Xiuyuan Lu, Yuki Hosono, Shigenari Ishizuka, Masamichi Nagae, Eri Ishikawa, Daisuke Motooka, Yuki Ozaki, Nicolas Sax, Ryo Shinnakasu, Takeshi Inoue, Taishi Onodera, Takayuki Matsumura, Masaharu Shinkai, Takashi Sato, Shota Nakamura, Shunsuke Mori, Teru Kanda, Emi E. Nakayama, Tatsuo Shioda, Tomohiro Kurosaki, Hisashi Arase, Kazuo Yamashita, Yoshimasa Takahashi, Sho Yamasaki

## Abstract

T cells play pivotal roles in protective immunity against SARS-CoV-2 infection. Follicular helper T (Tfh) cells mediate the production of antigen-specific antibodies; however, T cell receptor (TCR) clonotypes used by SARS-CoV-2-specific Tfh cells have not been well characterized. Here, we first identified and crystallized public TCR of Tfh clonotypes that are shared and expanded in unhospitalized COVID-19-recovered patients. These clonotypes preferentially recognized SARS-CoV-2 spike (S) protein epitopes which are conserved among emerging SARS-CoV-2 variants. These clonotypes did not react with S proteins derived from common cold human coronaviruses, but cross-reacted with symbiotic bacteria, which might confer the publicity. Among SARS-CoV-2 S epitopes, S_864-882_, presented by frequent HLA-DR alleles, could activate multiple public Tfh clonotypes in COVID-19-recovered patients. Furthermore, S_864-882_-loaded HLA tetramer preferentially bound to CD4^+^ T cells expressing CXCR5. In this study, we identified and crystallized public TCR for SARS-CoV-2 that may contribute to the prevention of COVID-19 aggravation.

## Introduction

Severe acute respiratory syndrome coronavirus 2 (SARS-CoV-2) has caused a worldwide pandemic with emerging mutations and continuous development of robust vaccine is imperative. Neutralizing antibodies are a major component of a protective immunity against SARS-CoV-2 (Tay et al., 2020), and thus information about virus-specific CD4^+^ T cells that help B cells to produce antigen-specific antibodies is needed. Among CD4^+^ T cells, follicular helper T (Tfh) cells mediate germinal center B cells maturation and production of high affinity antibodies (Crotty, 2019). Some deceased COVID-19 patients exhibited incomplete Tfh cell development (Kaneko et al., 2020), suggesting that efficient Tfh induction is a key to protective immunity against SARS-CoV-2 (Gong et al., 2020; Zhang et al., 2020). Furthermore, the characterization of SARS-CoV-2-specific Tfh cells bearing common TCR shared between different individuals will help us to understand and predict host responses during infection. However, such public Tfh clonotypes specific for SARS-CoV-2 have not yet been reported.

Given that V(D)J recombination is a random process, presence of shared TCR between individuals is generally thought to be a rare event (Soto et al., 2020; Venturi et al., 2008). Common antigens are one of the reasons that maintain such public clonotypes in the periphery (Li et al., 2012; Venturi et al., 2008). Recently, some T cells recognizing common cold human coronaviruses (HCoVs) are reported to cross-react with SARS-CoV-2 (Braun et al., 2020; Mateus et al., 2020; Woldemeskel et al., 2020), whereas SARS-CoV-2-specific T cells that do not react with HCoVs are frequently detected (Bacher et al., 2020; Mateus et al., 2020). Currently, other cross-reactive antigens recognized by SARS-CoV-2-specific T cells have not been well examined.

In this study, using single cell-based paired TCR analysis together with a rapid TCR reconstitution and epitope determination platform, we identified 1) public TCRs of Tfh cells specific for SARS-CoV-2 expanded in convalescent patients, 2) their epitope peptides and restricting HLA alleles, and 3) their cross-reactive proteins from symbiotic bacteria.

## Results

### Identification of SARS-CoV-2 spike protein-specific Tfh clones in convalescent COVID-19 patients

We first analyzed SARS-CoV-2-specific T cell subsets and their clonotypes using a single cell-based RNA sequencing platform. Peripheral blood mononuclear cells (PBMCs) isolated from healthy donors and convalescent COVID-19 patients (Table S1 and S2) were stimulated with antigens derived from SARS-CoV-2, including inactivated virus, recombinant S protein, overlapped peptide pools derived from S protein (S peptide pool) or membrane (M) and nucleocapsid (N) proteins (MN peptide pool). Activation marker-positive T cells were sorted and analyzed for their T cell receptor (TCR) sequences together with RNA expression by single cell TCR-seq and RNA-seq analyses (Figure S1). UMAP embedding and clustering allowed identification of a cluster (#8) consisting of CD4^+^ cells expressing Tfh-related genes, such as *CD200*, *PDCD1*, *ICOS*, *CXCL13*, *CD40LG* and *CXCR5*, suggesting that cluster #8 includes circulating Tfh cells (Crotty, 2019; Schmitt et al., 2014) (Figure 1A and 1B). Within this Tfh cluster, we could identify 1735 TCRs (Table S3). Furthermore, in the S-reactive Tfh cluster, we detected clones bearing TCRαβ pairs shared between different patients, Ts-017 and Ts-018, who had antibodies against receptor binding domain (RBD) of S protein and neutralizing antibodies (Table S1). Considering the cell number of single cell TCR-seq, these are likely to be frequent public TCRs and we designated them as TCR-017 and TCR-018, respectively. TCR-017 and -018 had the same Vα, Jα, Vβ and Jβ usages and their complementarity-determining region 3 (CDR3) sequences were identical except for one amino acid in CDR3β (Figure 1C and 1D). TCR-018 was detected in patient Ts-018 as three different barcoded clones within S protein/peptides-reactive Tfh cluster (Table S3).

**Figure 1.**
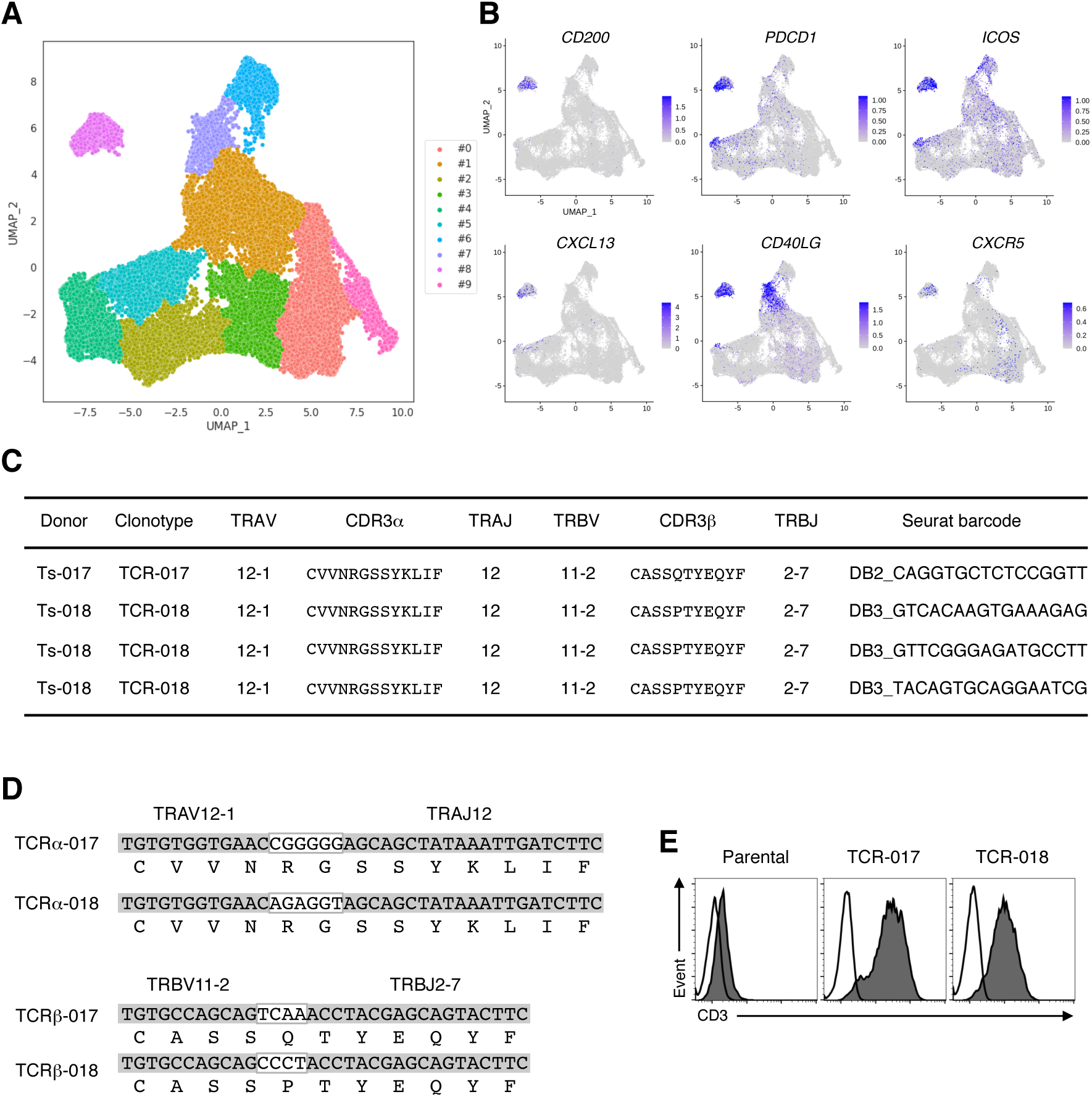
Shared TCR expressed by SARS-CoV-2-reactive Tfh cells from convalescent patients. (A) UMAP projection of T cells in single-cell analysis of PBMCs identified a cluster representing Tfh cells (cluster #8). Each dot corresponds to a single cell and was colored according to the cluster. (B) The expressions of canonical Tfh cell markers, CD200, PDCD1, ICOS, CXCL13, CD40LG and CXCR5 were shown as heatmaps in the UMAP plot. (C) Usages of variable (V) and Joint (J) gene segments and CDR3 sequences of α and β chains of TCR-017 and TCR-018. For TCR-018, three different barcoded clones having the same nucleotide sequences detected in Ts-018 were shown. (D) Alignment of CDR3α and CDR3β nucleotide and amino acid sequences of TCR-017 and TCR-018. Nucleotide sequences in white boxes indicate N-nucleotides. (E) Expression of paired TCRs on reporter cells indicated by surface CD3 expression. Closed, anti-mouse CD3 mAb; open, isotype control antibody. See also Figure S1 and Table S1-S3.

### Tfh clones derived from different patients recognized an identical S epitope restricted by the same HLAs

To examine the antigen specificity of TCR-017 and -018, the respective TCRα and β chains were reconstituted in a TCR-deficient T cell hybridoma bearing an NFAT-GFP reporter (Figure 1E). TCR-transfectants were stimulated with SARS-CoV-2 antigens in the presence of transformed B cells derived from the same patients as autologous antigen-presenting cells (APCs). Cells expressing TCR-017 and -018 responded to recombinant S protein and S peptide pool, but not to MN peptide pool (Figure 2A), which is consistent with the initial antigen specificity revealed in single cell analysis (Table S3). Among two halves of pooled S peptides, #1 (S_1-643_) and #2 (S_633-1273_), only #2 pool reacted with both TCRs (Figure 2B). However, another SARS-CoV-2 S peptide pool (S_304-338, 421-475, 492-519, 683-707, 741-770, 785-802, 885-1273_) did not activate either clone (Figure S2A and Figure 2B). The above results suggest that the antigen epitope(s) of TCR-017 and -018 is located in regions 633-682, 708-740, 771-784 or 803-884 of S protein (Figure S2A).

**Figure 2.**
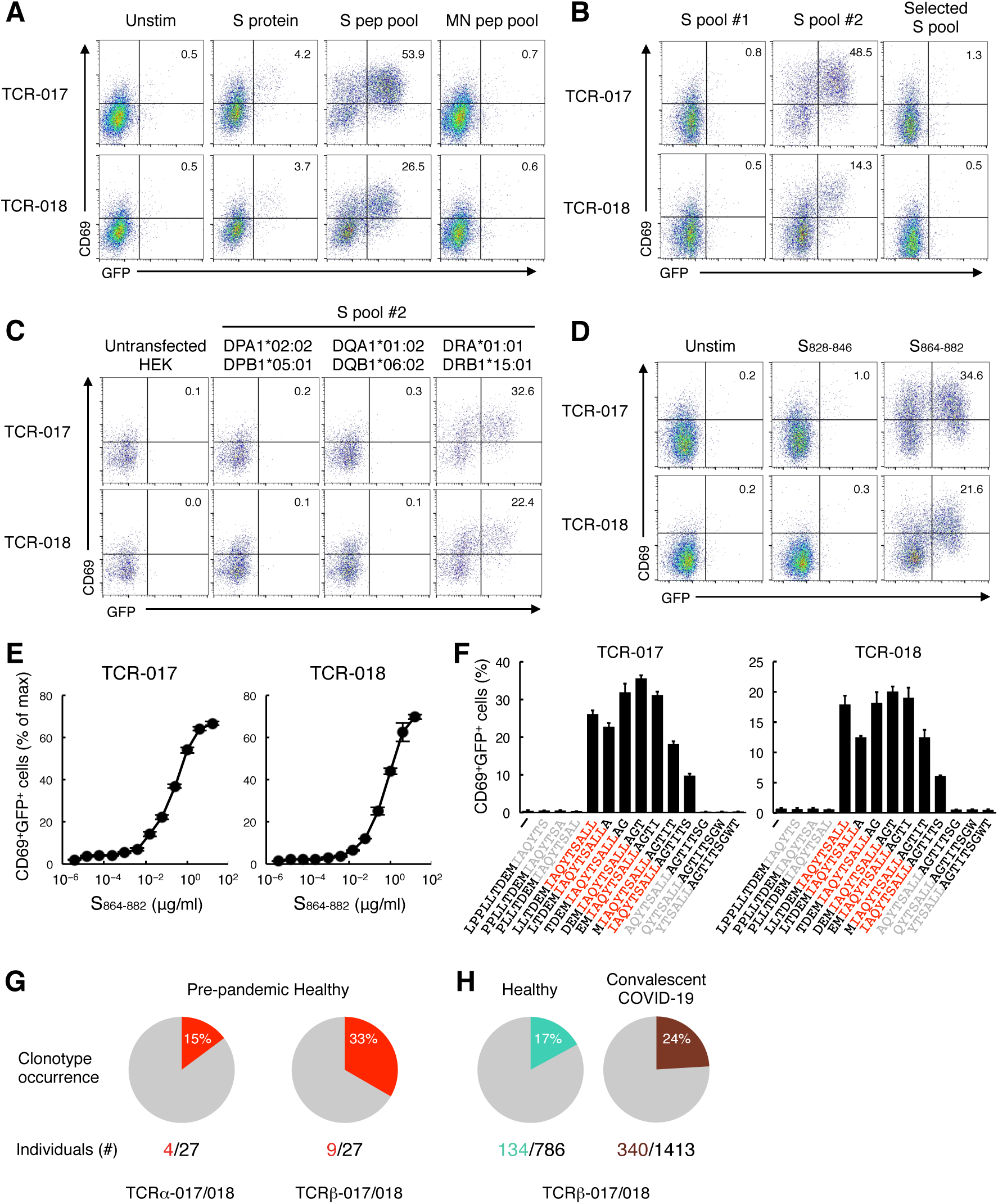
TCR-017 and -018 recognize the same core epitope, S_870-878_, presented on DRB1*15:01. (A) Cells expressing TCR-017 or TCR-018 were stimulated with 10 μg/ml recombinant S protein, 1 μg/ml S protein peptide pool (S pep pool) or 1 μg/ml M and N protein peptide pool (MN pep pool) in the presence of autologous APCs and analyzed for GFP and CD69 expressions. (B-D) Cells expressing TCR-017 or TCR-018 were stimulated with 0.3 μg/ml S peptide pool #1, #2 or Selected S pool (S_304-338, 421-475, 492-519, 683-707, 741-770, 785-802, 885-1273_) (B), 1 μg/ml S peptide pool #2 in the presence of HEK293T cells expressing indicated HLAs (C), or 1 μg/ml single peptides S_828-846_ or S_864-882_ (D). (E) Sensitivity of TCR-017 and TCR-018 to the epitope. TCR transfectants were stimulated with indicated amount of S_864-882_ and APCs from Ts-018. Activities were shown as percentages to the maximum responses induced by plate-coated anti-CD3 mAb. Data are shown in mean ± SD of triplicates. (F) Core epitope recognized by TCR-017 and TCR-018 (shown in red). Cells were stimulated with 1 μg/ml of serial overlapped 15-mer peptides covering S_861-887_ region. Data are shown in mean ± SD of triplicates. (G and H) Occurrence of TCRα-017/018 (CDR3: CVVNRGSSYKLIF) and TCRβ-017/018 (CDR3: CASS[Q or P]TYEQYF) in a pre-pandemic healthy cohort from Japan (n = 27) (G), and in cohorts of healthy donors (n = 786) and convalescent COVID-19 patients (n = 1,413) (H). Numbers of individuals possessing the TCRs among cohorts were indicated. All data are representatives of at least two independent experiments. See also Figure S2 and Table S3.

To further reduce the number of epitope candidates, we next determined the HLA alleles that restrict TCR-017 and -018. Heterologous APCs from patient Ts-017 could activate TCR-018, and *vice versa*, demonstrating that these APCs are interchangeable (Figure S2B). Thus, the shared HLA class II alleles, DRA-DRB1*15:01, DQA1*01:02-DQB1*06:02 or DPA1*02:02-DPB1*05:01 (shown in red in Table S2), are likely to restrict this recognition. Since DRB1*15:01 and DQB1*06:02 are genetically linked and would not be segregated from each other (Begovich et al., 1992), these HLA alleles were separately transfected into HEK293T cells to examine their ability to present S peptides to TCR-017 and -018. Only cells transfected with DRA and DRB1*15:01 could activate both clones in the presence of S peptide pool #2 (Figure 2C), suggesting that the reactivity is determined by DRB1*15:01, as DRA is monomorphic. Two epitopes, S_828-846_ and S_864-882_, were predicted as strong binders to DRA-DRB1*15:01 using NetMHC server software (Reynisson et al., 2020) within the candidate regions described above (Figure S2A). Among them, S_864-882_ was identified as the epitope for both clones (Figure 2D). Judging from their dose-response curves, relative affinities of TCR-017 and -018 to this peptide appeared comparable (Figure 2E). Furthermore, antigenic activities of serial overlapping peptides indicated that S_870-878_ is a minimum epitope recognized by these TCRs (Figure 2F). Taken together, both TCR-017 and -018 recognized the same epitope on the same HLA with similar affinities, indicating that TCR-017 and -018 are considered as one clonotype, clonotype-017/018, shared by different donors.

Among other HLAs predicted to strongly bind S_864-882_ (Reynisson et al., 2020), we found that DRB1*15:02/*15:03/*15:04, but not DRB1*16:02 and DQA1*01:01-DQB1*05:01, were restricting HLA that present this epitope to activate clonotype-017/018 (Figure S2C and S2D).

### Public clonotypes significantly expanded in recovered-COVID-19 patients preferentially recognize conserved S epitopes

Consistent with the high frequency of DRB1*15:02 (10.6%) and *15:01 (7.7%) in the Japanese population (http:/hla.or.jp), bulk TCR sequencing of pre-pandemic Japanese donors revealed that TCRα and β chains of clonotype-017/018 were broadly shared by 14.8% and 33.3%, respectively, of the population (Figure 2G). As DRB1*15 are not minor alleles worldwide (Gonzalez-Galarza et al., 2020), we analyzed TCR databases of healthy and convalescent COVID-19 cohorts containing donors of various ethnicities (Emerson et al., 2017; Nolan et al., 2020) and found that 17% and 24%, respectively, possessed TCRβ-017/018 (Figure 2H). These results suggest that clonotype-017/018 is a public clonotype in multiple ethnicities.

We therefore examined whether any of 1735 paired TCRs detected in our sample pool are found in these cohorts. 855 out of 1735 were revealed to be public Tfh TCRs (Table S3), indicating that TCR-017/018 is not an exceptional clonotype. Among them, we identified 10 Tfh clonotypes which were significantly expanded in recovered COVID-19 patients compared with the healthy cohort (Figure 3A, column J, asterisks). Of note, a half of such clonotypes were indeed detected in multiple patients in our sample pool by single cell TCR sequencing (Figure 3A, column A, #), and TCR-017/018 was the fifth most significantly expanded clonotype (Figure 3A, column A and J).

**Figure 3.**
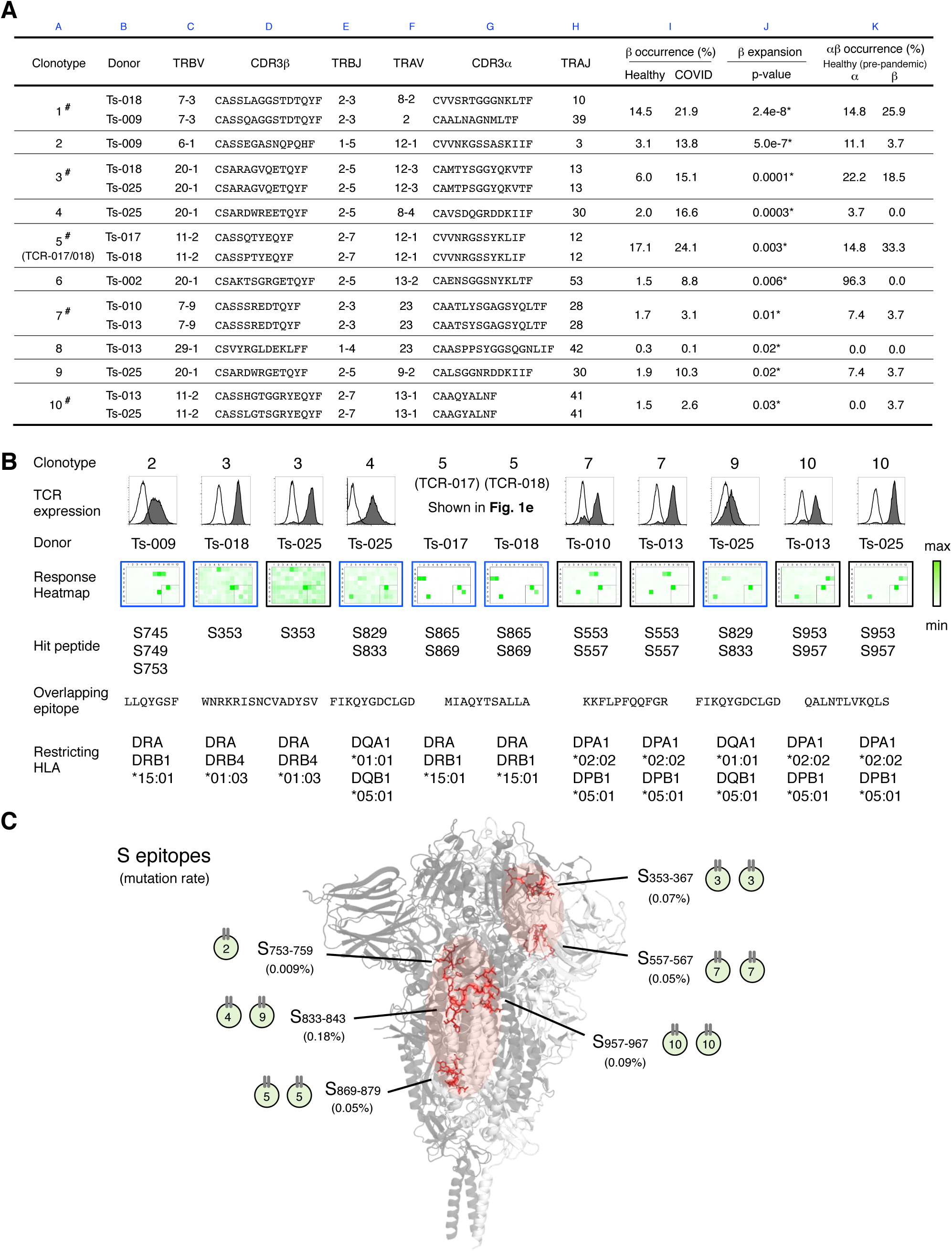
Determination of epitopes of public Tfh clonotypes expanded in COVID-19 patients within conserved S regions with low mutation rates. (A) Characterization of public clonotypes. Clonotypes detected in multiple donors from our sample pool are shown as # (column A). Clonotype occurrences were calculated based on a cohort of pre-pandemic healthy donors (n = 27) and public database including healthy donors (n = 786) and convalescent COVID-19 patients (n = 1,413) datasets (column K and I). For clonotypes that contain different TCRα or TCRβ sequences, occurrences are calculated using pooled sequences (column K and I). Clonotypes were listed following the order of significance of TCRβ expansion. *, Clonotypes expanded significantly in recovered patients (p < 0.05) (column J). (B) TCR pair of each clonotype listed in (A) was reconstituted in reporter cells and tested as described in Figure S3A in the presence of shared HLA-expressing HEK293T cells or autologous APCs. Expressions of paired TCRs on reporter cells indicated by surface CD3 expression were shown in histograms. Closed, TCR-pairs-reconstituted reporter cells; open, parental cells. GFP reporter activities are shown as heatmaps. Restricting HLAs were determined as described in Figure S3B-F, Table S2 and Methods. Clonotypes 1 and 6 did not respond to S peptide pools. TCRαβ of clonotype 8 was not expressed on the cell surface. (C) Locations of epitopes described in (B) are highlighted in red in 3D structure of SARS-CoV-2 S protein (PDB code: 6XR8). Frequencies of mutation (mutation rate) defined relative to Wuhan-Hu-1 were calculated based on the data from CoV-GLUE that enabled by data from GISAID. See also Figure S3 and Table S3.

To efficiently determine the epitopes of these clonotypes, we established a rapid epitope-determination platform (Figure S3A) and validated it using TCR-017/018 (Figure 3B, clonotype 5). By using this platform, we further determined the epitopes for other COVID-19-expanded Tfh clonotypes (Figure 3B). Restricting HLAs of these epitopes were determined using overlapped HLA alleles in the combination of transformed B cells from different individuals, and individual HLA transfectants (Table S2 and Figure S3B-S3F). These results indicate that the public Tfh clonotypes which significantly expanded in unhospitalized, recovered patients preferentially recognized S epitopes (Figure 3A and 3B). All identified epitopes were located in two major hotspots within the conserved trimer-forming interface of S protein with low mutation rates (Figure 3C)(Elbe and Buckland-Merrett, 2017; Singer et al., 2020). In addition, the donors of these clonotypes had anti-RBD and neutralizing antibodies (Table S1). These results suggest that public Tfh clonotypes specific for conserved S epitopes have expanded during SARS-CoV-2 infection and presumably contributed to the recovery from mild symptoms (Emerson et al., 2017; Nolan et al., 2020).

### Crystal structure revealed sequence flexibility of CDR3 that extends the publicity of SARS-CoV-2-specific clonotypes

We demonstrated that TCRβ-017/018 is highly public and expanded in the recovered patients (Figure 3A). Since the nonidentical amino acid, Q94 and P94 of TCRβ-017 and -018, respectively, have distinct characteristics, we suspected that any amino acid at this position allows epitope recognition. Indeed, any of 17 substitutions, except for D94, retained the reactivity to S_864-882_ epitope on DRA-DRB1*15:01 (Figure 4A). Indeed, crystallographic analysis of TCR-017 complex demonstrated that the residue at position 94 in CDR3β is located away from CDR3α (Figure 4B and 4C) and thus unlikely to contact with peptide-MHC. In addition, C90, A91, S92 and S93 were more distant from CDR3α (Figure 4C), suggesting that this TCRβ may mainly utilize a C-terminal half of CDR3β for recognition and allow diverse TCR sequence variants or even other TRBVs.

**Figure 4.**
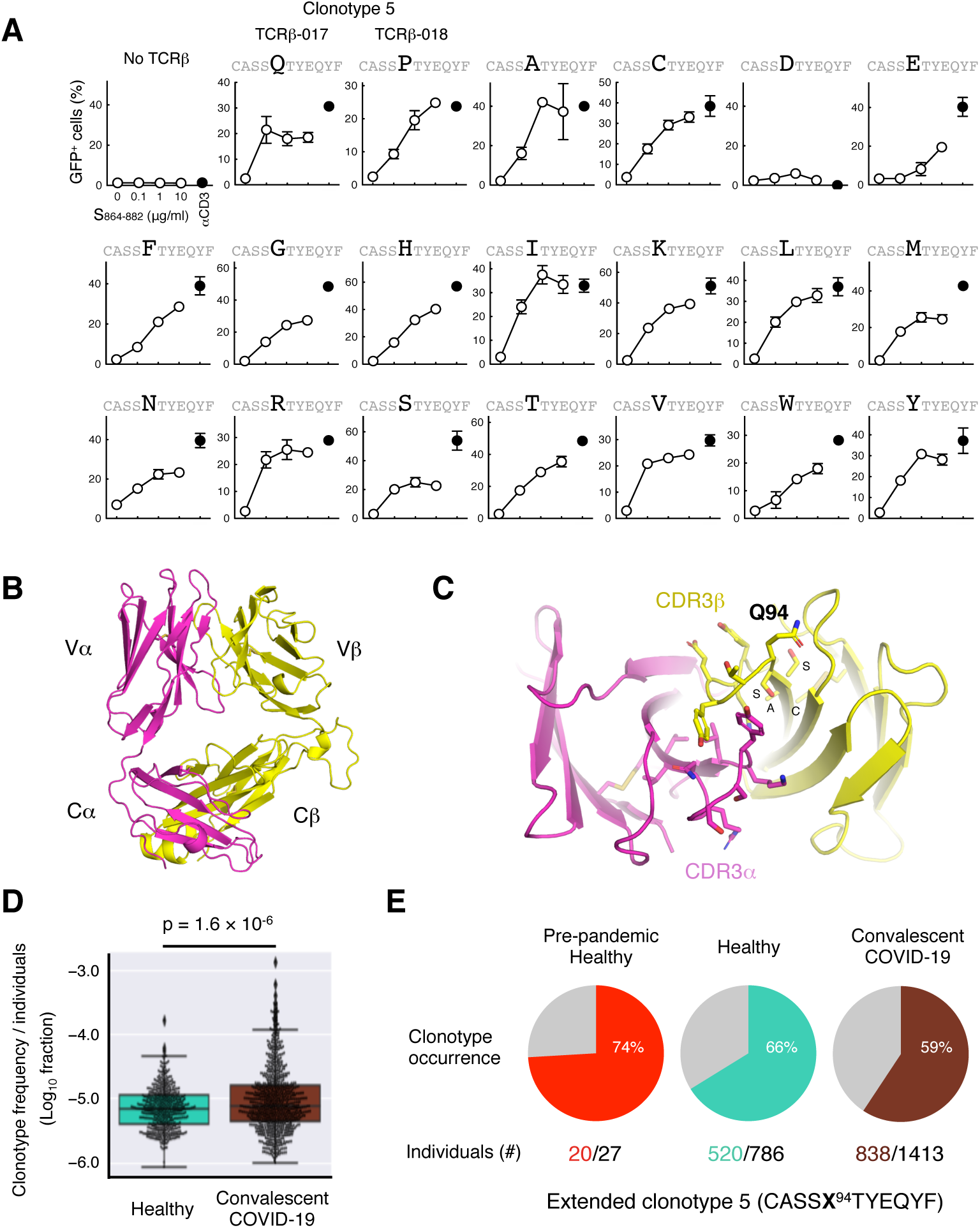
Structural basis of peptide recognition by public clonotype 5 (017/018). (A) TCRβ-017, -018 or mutants at position 94 were co-expressed with TCRα-017/018 and human CD4 in reporter cells. Cells were kept unstimulated or stimulated with 1 μg/ml of peptide S_864-882_ or anti-CD3 antibody in the presence of APCs from Ts-018. Percentages of GFP^+^ cells in the CD3^+^ populations are shown. Data are shown in mean ± SD of duplicates. (B and C) Crystal structure of TCRαβ heterodimer of clonotype 5 (TCR-017). Whole view (B) and close-up view of CDR3 domains (C) are shown. Q94, C90, A91, S92 and S93 within CDR3β are indicated. (D) Frequencies of functional antigen-reactive TCRβ chains harboring acceptable 17 substitutions at position 94 except for D94 of TCRβ-017/018 determined in (A) in the healthy donors and convalescent COVID-19 patients. Horizontal line, median; bottom/top, first/third quartile; whiskers, 1.5 times the interquartile range. (E) Occurrences of extended antigen-reactive TCRβ chains harboring acceptable 17 substitutions at position 94 (shown as X) except for D94 of TCRβ-017/018 determined in (A) in the individuals in a pre-pandemic Japanese healthy cohort, a healthy cohort and a convalescent COVID-19 cohort described in Figure 2G and 2H. Numbers of individuals possessing this TCRβ among cohorts are indicated. See also Table S7.

This promiscuity within CDR3β sequence should expand the pool of this public clonotype, which gives us a more precise expansion rate in recovered patients. Indeed, the expansion of the extended clonotype in the COVID-19-recovered cohort became much more significant (p = 1.6 × 10^−6^) (Figure 4D) than the value shown in Figure 3A (p = 0.003). Furthermore, its occurrence became more than half in all three cohorts (Figure 4E), suggesting that the extended clonotype 5 is broadly shared among a large population.

### Public Tfh clonotypes did not cross-react with HCoVs but with symbiotic bacteria

We next asked how these clonotypes, particularly clonotype 5, acquired publicity. Cross-reactivity with the common cold coronaviruses is one of the possible mechanisms (Braun et al., 2020; Grifoni et al., 2020; Mateus et al., 2020). However, none of the clonotypes identified in this study including TCR-017/018 reacted with S proteins derived from common cold human coronavirus (HCoV) strains (Figure 3B and Figure S3A, S4A and S4B). Indeed, these HCoVs do not have identical sequences to SARS-CoV-2 epitopes determined in Figure 3B (Figure 5A). Furthermore, peptides in the corresponding regions derived from these coronaviruses including severe acute respiratory syndrome coronavirus (SARS-CoV) and Middle East respiratory syndrome-related coronavirus (MERS-CoV) were not recognized by TCR-017/018 (Figure S4C and S4D). Thus, these clonotypes are unlikely to acquire publicity by cross-reacting with reported human coronaviruses present before the outbreak of COVID-19. This raises a hypothesis that some other common antigen(s) may confer publicity on these clonotypes. In line with this, BLAST homology search of S_870-878_ core epitope revealed that oral commensal bacteria, *Selenomonas noxia*, express a protein, multidrug and toxic compound extrusion (MATE) family efflux transporter, that includes the same epitope with S_870-879_, IAQYTSALLA (Figure 5A). Indeed, the synthetic peptides (MATE_241-260_ and _242-256_) were recognized by both TCR-017 and -018 on DRB1*15:01 (Figure 5B). In addition, if any amino acids within this S epitope are replaceable, further cross-reactive proteins might be found. To identify such acceptable substitutions, we performed Alanine/Glycine scanning of S_870-878_ peptides and found that seven single substitutions, except for positions Q872 and T874, retained antigenicity of the epitope (Figure 5C). Among these mutated epitopes, the sequence of L877A (IAQYTSA<u>A</u>L) was found in a protein expressed by various bacteria, histidine ammonia-lyase (HAL, EC 4.3.1.3)(Table S4 and Figure 5A).

**Figure 5.**
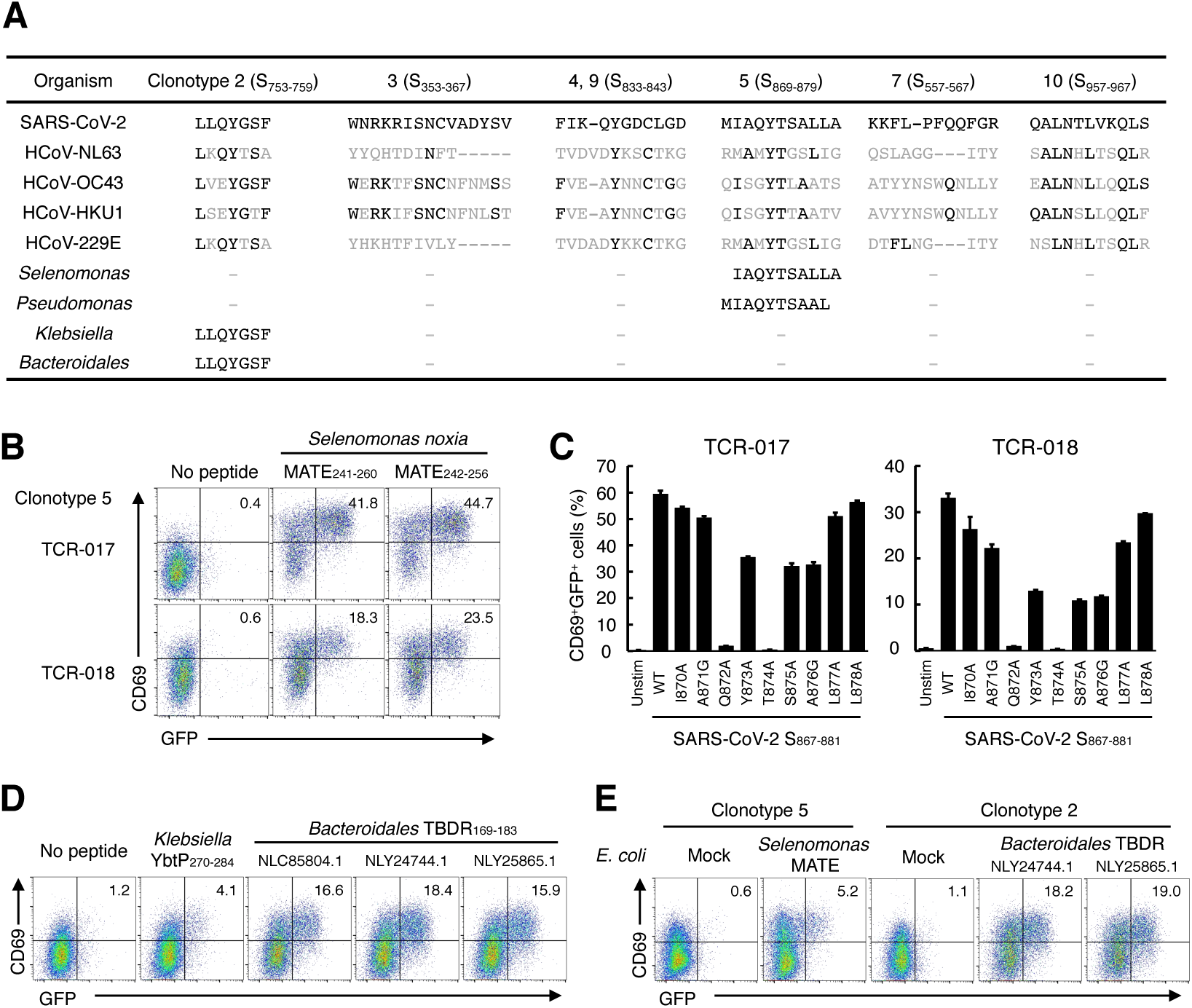
Cross-reactivity of SARS-CoV-2-specific public Tfh clonotypes. (A) Alignment of cross-reacting epitopes. Similar amino acids to SARS-CoV-2 epitopes found in HCoVs and microbes are shown in black. Details of the microbes and full list of homologous search are shown in Table S4 and S5. (B) Cells expressing clonotype 5 (TCR-017 or TCR-018) were stimulated with 1 μg/ml of peptides derived from MATE family efflux transporter of *Selenomonas noxia*. (C) Cells expressing clonotype 5 (TCR-017 or TCR-018) were stimulated with 1 μg/ml of S_867-881_ peptides containing indicated mutations in the presence of APCs from Ts-018. Data are shown in mean ± SD of triplicates. (D) Cells expressing TCR from clonotype 2 were stimulated with 1 μg/ml of peptides derived from *Klebsiella pneumoniae* yersiniabactin ABC transporter ATP-binding/permease protein (YbtP) or *Bacteroidales* bacterium TonB-dependent receptor (TBDR). Accession numbers of TBDRs from different *Bacteroidales* bacteria are shown. (E) Cells expressing TCRs from clonotype 5 (TCR-017) or clonotype 2 were stimulated with heat-killed *E. coli* transformed with the expression vectors for indicated microbial proteins in the presence of monocyte-derived dendritic cells expressing DRB1*15:01. See also Figure S4 and Table S4 and S5.

Other 9 public Tfh clonotypes significantly expanded in patients were also unresponsive to HCoVs (Figure 3B). Instead, homology searches for the epitopes of these clonotypes (Figure 3B) revealed that S_753-759_, an epitope of clonotype 2, was contained in proteins of various organisms including symbiotic bacteria, such as *Bacteroidales* bacteria and *Klebsiella pneumoniae* (Atarashi et al., 2017; Sefik et al., 2015) (Table S5 and Figure 5A). Indeed, these bacterial peptides, such as TonB-dependent receptor (TBDR)_169-183_ and yersiniabactin ABC transporter ATP-binding/permease protein (YbtP)_270-284_, respectively, could activate T cells reconstituted with clonotype 2 TCR (Figure 5D). Furthermore, *E. coli* transformed with the expression vectors for these proteins could activate public clonotypes in the presence of HLA-matched dendritic cells (Figure 5E), confirming that cross-reactive antigenic epitopes are processed and presented by APCs. These observations suggest that continuous priming via symbiotic bacteria might explain the high frequency of the public clonotypes worldwide (Figure 4E). Further cohort studies will clarify the etiological relationships between these bacteria and resistance against SARS-CoV-2 infection.

### S_864-882_ could activate multiple public Tfh clonotypes

Among 1735 TCRs derived from Tfh cluster (Table S3), we identified another clonotype from patient Ts-017 that recognizes the exact same epitope, S_864-882_, restricted by DRB1*15:01, despite it had distinct V/J usages and CDR3 sequences/lengths from TCR-017/018 (Figure S5). This clonotype was also public as it was detected in cohorts (Table S3, Seurat barcode, DB2_CAAGAAAGTGCGGTAA). Thus, this epitope, S_864-882_, could activate multiple public Tfh clonotypes even in the same patient.

To further examine the ability of this epitope to activate multiple Tfh clonotypes, PBMCs from additional convalescent/healthy donors possessing DRB1*15:01/*15:02 were stimulated with S_864-882_ *in vitro*. After epitope-induced expansion for 10 days, CD4^+^ T cells were subjected to single cell-TCR- and RNA-seq to determine the antigen-responsive clonotypes (Figure 6A and Table S6). As a result, we newly identified 104 public Tfh clonotypes that responded to S_864-882_ after filtering by two criteria; 1) expanded during stimulation based on single cell sequencing (> 0.1% of total CD4^+^ T cells) and 2) detected in cohort databases (Table S6, column K). Indeed, bulk TCR sequencing demonstrated that both α and β chains of all 104 clonotypes were expanded after S_864-882_ stimulation (Table S6, column I). For most of these clonotypes, Tfh characteristics were observed even after *in vitro* stimulation (Table S6, column J). These results indicate that S_864-882_ epitope is capable of expanding multiple public Tfh clones in DRB1*15:01/02 donors.

**Figure 6.**
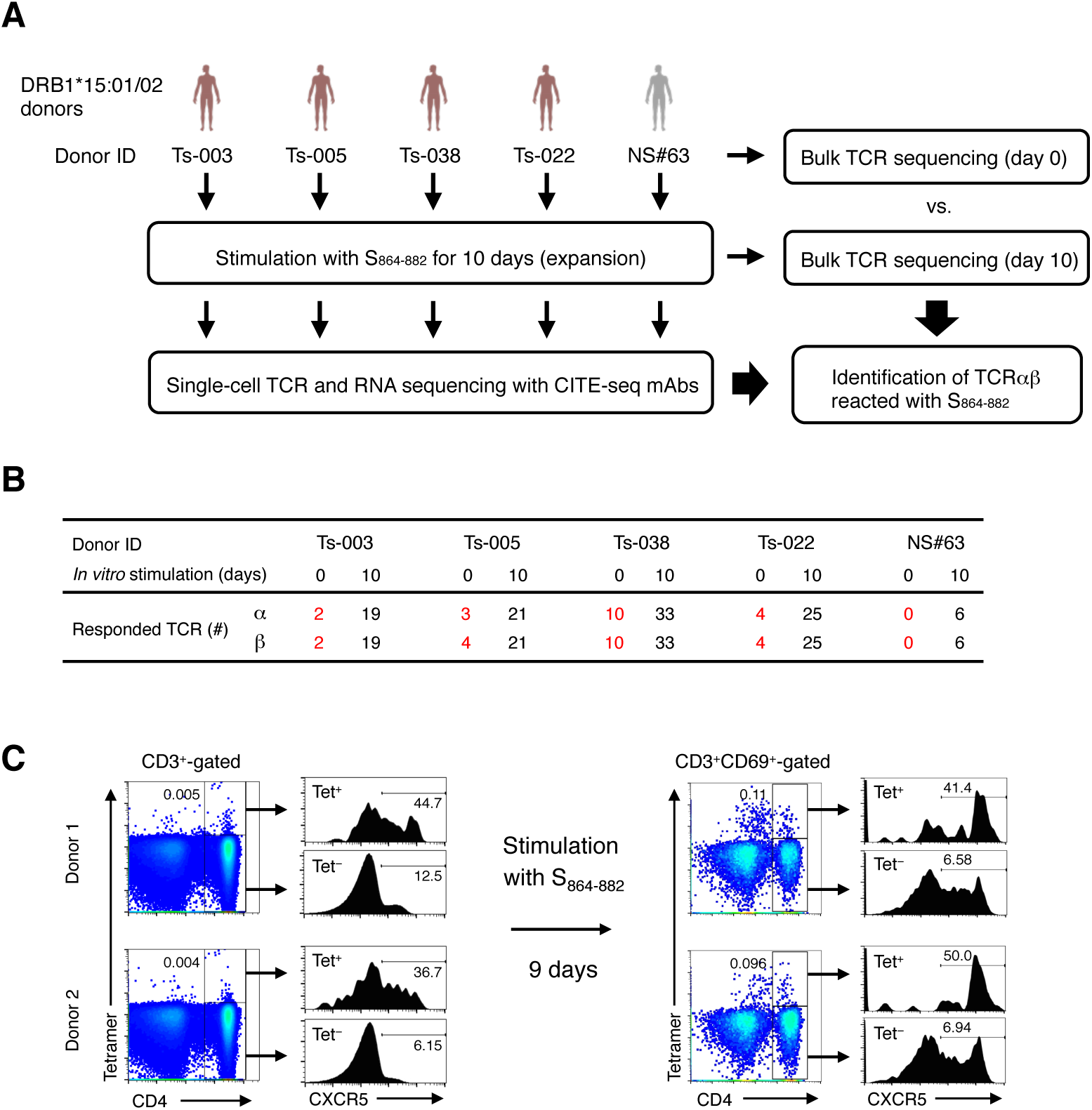
Characterization of S_864-882_-reactive T cells. (A) Workflow of the analysis for the S_864-882_-induced T cell expansion. PBMCs from convalescent COVID-19 patients (red) / healthy (grey) donors possessing DRB1*15:01 or *15:02 were analyzed by bulk TCR-seq, or stimulated with 1 μg/ml of S_864-882_ peptide for 10 days before CD4^+^ T cells were sorted and analyzed by single cell TCR-and RNA-seq as well as bulk TCR-seq. (B) Numbers of TCRα and β of expanded clonotypes (proportion > 0.1% in single cell TCR-seq) detected in bulk TCR-seq described in Table S6 are shown. (C) PBMCs from healthy donors possessing DRB1*15:01 were stained with S_867-881_-loaded DRA-DRB1*15:01 tetramer along with anti-CD4 and anti-CXCR5 antibodies before or after S_864-882_ stimulation followed by flowcytometric analysis. Percentages of CXCR5^+^ cells within tetramer-positive (Tet^+^) and -negative (Tet^-^) CD4^+^ cells are shown. All data are representative of at least two independent experiments. See also Figure S5 and S6 and Table S6.

To examine the presence of these S_864-882_-reactive clones in the periphery of donors before *in vitro* expansion, we assembled all expanded TCRαβ pairs with bulk TCR sequences of unstimulated peripheral T cells (Figure 6A, day 0). Approximately 20% of S_864-882_-reactive TCRα and β chains detected at day 10 (Table S6) were present in the ‘day 0’ bulk TCR sequences from all recovered patients possessing anti-S protein antibodies (Table S1), whereas none of such S_864-882_-reactive TCRs were detectable in healthy control before stimulation (Figure 6B). This data provides an implication that some S_864-882_-reactive Tfh clones have been expanded in recovered patients during SARS-CoV-2 infection and maintained as a memory pool after infection.

To directly detect S_864-882_-reacting T cells in PBMCs, we generated DRA-DRB1*15:01 tetramer loaded with this epitope, which was confirmed to bind to recombinant soluble TCR-017αβ *in vitro* (Figure S6A and S6B). In healthy donors possessing DRB1*15:01, tetramer-positive CD4^+^ T cells (approximately 0.005% of T cells) expressed higher CXCR5 than tetramer-negative CD4^+^ cells (Figure 6C). After *in vitro* activation by S_864-882_ peptide, the percentages of tetramer-positive CD4^+^ T cells were markedly increased and CXCR5^+^ population was further accumulated (Figure 6C), suggesting that S_864-882_ peptide is capable of generating activated CD4^+^ T cells with Tfh profiles.

## Discussion

Recently, public TCR for SARS-CoV-2 has been extensively studied (Shomuradova et al., 2020). In this study, we report the first identification and structural determination of SARS-CoV-2-specific public TCR of CD4^+^ T cells. Furthermore, we identified multiple public Tfh clonotypes that were significantly expanded in COVID-19-recovered patients by preferentially recognizing conserved S epitopes.

These observations suggest that the individuals possessing such public Tfh clonotypes could efficiently respond to S protein and produce antibodies against the cognate antigen, which may contribute to the prevention of disease aggravation and subsequent recovery (Table S1)(Emerson et al., 2017; Nolan et al., 2020). As these public clonotypes were widely detected in multiple ethnicities, the precise monitoring of the frequencies of public Tfh clonotypes using tetramer or clonotypic mAb could provide a novel option for prognosis prediction. Furthermore, such common S epitopes are candidates for next-generation vaccine that promote Tfh responses against SARS-CoV-2 and prevent aggravation.

The current study still has several limitations. 1) We analyzed only PBMCs. Although circulating Tfh cells in the periphery can reflect Tfh cells in germinal center (Crotty, 2019; Heit et al., 2017; Hill et al., 2019; Locci et al., 2013; Schmitt et al., 2014), the analysis of tissue resident Tfh cells of biopsy samples is ideal to fully characterize Tfh clonotype. 2) Although the present study suggests that these Tfh clonotypes contribute to amelioration of COVID-19 in patients recovered from mild symptoms, additional studies on patients with severe symptoms are helpful for further aspects. 3) The restricting HLA of the public clonotypes expanded in the patients were DRA-DRB1*15:01 (8.8%), DRA-DRB1*15:02 (2.5%), DRA-DRB1*15:03 (2.0%), DRA-DRB1*15:04 (0.02%), DRA-DRB4*01:03 (33.9% in local)(González-Quezada et al., 2019), DPA1*01:03 (60.7%)-DPB1*04:02 (16.5%), and DPA1*02:02 (24.7%)-DPB1*05:01 (14.8%), which cover a large part of, but not all, population worldwide (Gonzalez-Galarza et al., 2020).

Notwithstanding, the current study proposes that combination of single cell paired TCR sequencing and public database is a potent tool to identify public clonotypes which actually responded to SARS-CoV-2 in patients. The rapid reconstitution and determination platform enables us to identify epitopes and responsible HLA simultaneously. As the identified T cell epitopes are conserved among emerging SARS-CoV-2 mutants (Elbe and Buckland-Merrett, 2017; Singer et al., 2020) (Figure 3C), these peptides could become candidates for a robust vaccine against mutants, such as B.1.1.7 (UK), B.1.351 (South Africa), P.1 (Brazil) and presumably other future variants (Faria et al., 2021; Rambaut et al., 2020; Tegally et al., 2020). Furthermore, the identification of cross-reacting symbiotic bacteria in this study would provide novel prognosis prediction and therapeutic options. To extend a peptide pool for the vaccine development, identification of additional public clonotypes and their epitopes is necessary and now under investigation.

## Acknowledgements

We thank T. Ito, K. Toyonaga, Y. Adachi, S. Moriyama, Y. Harima and D. Azizah for experimental support; C. Schutt, W. Ise, J. B. Wing, D. Okuzaki, D. Standley, S. Teraguchi, H. Nakagami, S. Futami and T. Kobayashi for discussion. This research was supported by AMED (19fk0108161h0001 (HA), JP20fk0108265, JP20nf0101623, JP20gm0910010, JP20ak0101070 and JP20fk0108075 (SY)) and JSPS KAKENHI JP18H05279 (HA).

## Author Contributions

S.Y. conceptualized research; X.L., Y.H., S.I., E.I., M.N. and Y.O. did investigation; R.S., T.I., T.O., T.M., M.S., T.S., S.M., T.K., E.N., T.S., T.K., H.A. and Y.T. provided resources; D.M., N.S., S.N. M.N. and K.Y. did data curation; S.Y. supervised the research; X.L., Y.H. and S.Y. wrote the manuscript.

## Methods

### Human subjects

The Institutional Review Boards of Osaka University (approval number 898-4) and National Institute of Infectious Diseases (approval number 1137) approved blood draw protocols for convalescent COVID-19 and healthy individuals. The Institutional Review Board of nonprofit organization MINS (approval number 190210) approved the analyses of blood samples from healthy individuals in KOTAI Biotechnologies. The research was performed in accordance with all relevant guidelines and regulations. Written informed consent was obtained from all participants or designated healthcare surrogates if participants were unable to provide informed consent. Study enrollment criteria included subjects over 20 years old, regardless of disease severity and genders (Table S1). Prior to enrollment in this study, all COVID-19 donors were confirmed to be positive for SARS-CoV-2 by PCR using nasopharyngeal swab specimens. Blood from convalescent COVID-19 donors and healthy donors was obtained at Tokyo Shinagawa Hospital and Osaka University. Samples were de-identified prior to analyses. For PBMC preparation, whole blood was collected in heparin-coated tube and centrifuged to separate the cellular fraction and plasma, followed by density-gradient sedimentation. For neutralizing antibody assay, plasma samples from patients were heat-inactivated for 30 min at 56°C. Plasma diluted at 1:5 followed by two-fold serial dilutions was incubated with equal volume of solution containing 100 TCID_50_ of SARS-CoV-2 for 1 hour at 37°C and added to VeroE6/TMPRSS2 cells. After incubation for 5 days, the highest plasma dilution that protected 100% of cells from cytopathic effect was taken as the neutralization titer. Disease severity was defined as mild, moderate I, moderate II or severe in accordance with the Japanese COVID-19 clinical practice guideline ver. 2.2 (https://www.mhlw.go.jp/content/000650160.pdf).

### SARS-CoV-2

For the preparation of inactivated SARS-CoV-2, SARS-CoV-2 (KNG19-020) was kindly supplied by Dr. Tomohiko Takasaki (Kanagawa Prefectural Institute of Public Health). The virus was propagated in VeroE6/TMPRSS2 cells (JCRB1819) and purified by sucrose gradient centrifugation (Dent and Neuman, 2015). Concentrated virus was then inactivated by ultraviolet light (0.6 J/cm^2^).

### Antigens

Inactivated virus was used at 2.9 × 10^6^ viral particles/ml (approximately 1 μg S protein/ml) for stimulation. For the preparation of recombinant SARS-CoV-2 S protein, the ectodomain of S protein was cloned into a mammalian expression vector pCMV, with a foldon sequence followed by 9 × His-tag and Strep-tag at the C-terminus. Polybasic cleavage site (RRAR) of S protein was replaced by a single alanine, and K986P and V987P mutations were introduced to stabilize the conformation as previously described (Amanat et al., 2020). After transient transfection into Expi293F cells (Gibco), secreted protein was purified from the culture supernatant 4 days post transfection using TALON metal affinity resin (Clontech) and Amicon Ultra 10K filter device (Millipore). For the expression of S proteins of SARS-CoV-2 and HCoV-OC43, pME18S expression vectors for each protein were transfected into HEK293T cells. For the expression of bacterial proteins in *E. coli*, cDNA for MATE family efflux transporter, TonB-dependent receptor and TonB-dependent receptor plug domain-containing protein were cloned into pCold expression vector and transformed into *E. coli* BL21 competent cells Champion21 (SMBIO Technology). The protein expressions were induced by the addition of 0.5 or 1.0 mM isopropyl-β-D-thiogalactopyranoside (IPTG) at 37°C. Bacterial pellets were washed and resuspended with PBS and incubated at 95°C for 5 min for inactivation. The expressions of the proteins were confirmed by SDS-PAGE and CBB staining. PepMix SARS-CoV-2 (Spike Glycoprotein)(contains pool #1 and #2) was purchased from JPT Peptide Technologies. PepTivator SARS-CoV-2 for S, M and N protein were purchased from Miltenyi Biotec. Domains of SARS-CoV-2 S protein showed in Figure S2A was previously described (Wrapp et al., 2020). All of other peptides were from GenScript. 3D structure of SARS-CoV-2 S protein (PDB code: 6XR8) was depicted with program PyMOL ver. 2.0 (The PyMOL Molecular Graphics System, Version 2.0 Schrödinger, LLC.).

### In vitro stimulation of PBMCs

Cryopreserved PBMCs were thawed and washed with RPMI 1640 medium (Sigma) supplemented with 5% human AB serum (GeminiBio), Penicillin (Sigma), streptomycin (MP Biomedicals) and 2-mercaptoethanol (Nacalai Tesque). 5 × 10^5^ PBMCs were stimulated in the same medium with inactivated SARS-CoV-2, 1 or 10 μg/ml of recombinant S protein, 1 μg/ml of S peptide pool or 1 μg/ml of M + N (MN) peptide pool for 20 hours, followed by staining with anti-human CD3, CD69, CD137, CD154 and TotalSeq-C Hashtags antibodies. CD69^+^, CD137^+^ or CD154^+^ cells within CD3^+^-gated population were sorted by cell sorter SH800S (SONY) and used for single cell TCR and RNA sequencing analyses. For epitope-specific clonal expansion, 1-5 × 10^5^ PBMCs were stimulated with 1 μg/ml of S_864-882_ for 10 days and recombinant human IL-2 (1 ng/ml, Peprotech) was added at day 4 and day 7. CD4^+^ T cells were sorted and analyzed by single cell TCR and RNA sequencing.

### Single cell-based transcriptome and TCR repertoire analysis

Single cell capturing and library preparation were performed using following reagents; Chromium Next GEM Single Cell 5′ Library & Gel Bead Kit v1.1, 16rxns, PN-1000165; Chromium Next GEM Chip G Single Cell Kit, 48rxns, PN-1000120; Chromium Single Cell V(D)J Enrichment Kit, Human T Cell, 96 rxns, PN-1000005; Single Index Kit T Set A, 96 rxns, PN-1000213; Chromium Single Cell 5′ Feature Barcode Library Kit, 16 rxns, PN-1000080; Single Index Kit N Set A, 96 rxns, PN-1000212. Single cell suspension containing approximately 2 × 10^4^ cells were loaded into Chromium microfluidic chips to generate single-cell gel bead-in-emulsion using Chromium controller (10x Genomics) according to the manufacturer’s instructions. RNA from the barcoded cells for each sample was subsequently reverse-transcribed inside gel bead-in-emulsion using Veriti Thermal Cycler (Thermo Fisher Scientific), and all subsequent steps to generate single-cell libraries were performed according to the manufacturer’s protocol, with 14 cycles used for cDNA amplification. Then ∼50 ng of cDNA was used for gene expression library amplification by 14 cycles in parallel with cDNA enrichment and library construction for TCR libraries. Fragment size of the libraries was confirmed with Agilent 2100 Bioanalyzer (Agilent). Libraries were sequenced on Illumina NovaSeq 6000 as paired-end mode (read1: 28bp; read2: 91bp). The raw reads were processed by Cell Ranger 3.1.0 (10x Genomics). Gene expression-based clustering was performed using the Seurat R package (v3.1)(Hafemeister and Satija, 2019). Briefly, cells with a mitochondrial content above 10%, cells with less than 200 or more than 4,000 genes detected were considered outliers (dying cells, empty droplets and doublets, respectively) and filtered out. The Seurat SCTransform function was used for normalization, and data was integrated without performing batch-effect correction as all samples were processed simultaneously. Hashtag oligo demultiplexing was performed on CLR-normalized hashtag UMI counts, and clonotypes were matched to the gene expression data through their droplet barcodes, using Python scripts. Only cells assigned a single hashtag and a beta-chain clonotype were retained for downstream analyses.

### Bulk TCR sequencing and analysis

1-3 × 10^5^ PBMCs were lysed in QIAzol (Qiagen). Full-length cDNA was then synthesized using the SMARTer technology (Takara Bio), and the variable region of TCRα and β genes were amplified using TRAC/TRBC-specific primers. After sequencing of the variable region amplicons, each pair of reads was assigned a clonotype (defined as TR(A/B)V and TR(A/B)J gene, and the complementarity-determining region 3) using the MiXCR software (Bolotin et al., 2015). For each α/β clonotype, expansion was defined as the fraction of reads for that clone divided by the total number of reads for the α/β chain, respectively, and fractions were converted to log10 for plotting and statistical analyses.

### HLA class II typing

For HLA class II typing, genomic DNA samples were isolated from PBMCs using QIA DNA Mini Kit (Qiagen). AllType FASTplex NGS 11 Loci Flex Kit (ONE LAMBDA) was used to prepare DNA sequence libraries (DPA1, DPB1, DQA1, DQB1, DRB1 and DRB3/4/5) according to the manufacturer’s protocol. Sequencing was performed on Illumina MiSeq (Illumina). TypeStream Visual ver. 2.0 (ONE LAMBDA) was used to analyze the DNA sequences. For DRB1*15:01/02 typing, genomic DNA was amplified with a DRB1*15/16-specific forward primer (5’-CGT TTC CTG TGG CAG CCT AAG AGG-3’) and a DRB1*15-specific reverse primer (5’-CCG CGC CTG CTC CAG GAT-3’) followed by DNA sequencing.

### Antigen-presenting cells

Recombinant Epstein-Barr virus (EBV) was produced as previously reported (Kanda et al., 2015). Viral stocks were obtained by concentrating virus-containing culture supernatants by ultracentrifugation at 32,000 rpm for 1 hour. 3 × 10^5^ PBMCs were incubated with an aliquot of the viral stock for 1 hour at 37°C. The infected cells were cultured with RPMI 1640 medium supplemented with 20% fetal bovine serum (FBS, CAPRICORN SCIENTIFIC GmbH) containing 0.1 μg/ml cyclosporine A (CsA, Cayman Chemical). EBV-immortalized B lymphoblastoid cell lines were obtained after 3 weeks culture and used as APCs. To generate APCs expressing specific HLA, HEK293T cells were transfected with plasmids encoding HLA class II alleles as previously described (Jiang et al., 2013). EBV experiments were approved by the Minister of Education, Culture, Sports, Science and Technology (approval number 539) and the Institutional Review Boards of Osaka University (approval number 04658). To generate monocyte-derived dendritic cells, CD14^+^ cells were isolated from PBMCs using CD14 MicroBeads, human (Miltenyi Biotec), and cultured in RPMI 1640 medium supplemented with 10% FBS, 0.1 mM Non-Essential Amino Acid Solution (Gibco), 1mM Sodium Pyruvate (Gibco), 10 ng/ml of human GM-CSF (Peprotech) and IL-4 (Peprotech) for 6 days.

### TCR reconstitution and stimulation

TCRα and β chain cDNA sequences were synthesized and cloned into retroviral vectors pMX-IRES-rat CD2 and pMX-IRES-human CD8, respectively. Two vectors containing paired TCRα and β chains were transfected together into Phoenix packaging cells using PEI MAX (Polysciences). Supernatant containing retroviruses was used for infection into mouse T cell hybridoma with an NFAT-GFP reporter gene (Matsumoto et al., 2021) to reconstitute TCRαβ pairs. TCRβ mutants were constructed by site-directed mutagenesis. For antigen stimulation, TCR-reconstituted cells were co-cultured with stimulants in the presence of immortalized autologous B cells unless indicated otherwise. After 20 hours, T cell activation was assessed by GFP or CD69 expression.

### Rapid epitope-determination platform

For the preparation of pooled peptide matrix, 15-mer peptides with 11 amino acids overlap that cover the full length of S protein of SARS-CoV-2 were individually synthesized (GenScript). Peptides were dissolved in DMSO at 12 mg each peptide/ml and 8-12 peptides were mixed to create 75 different semi-pools so that the responsible epitopes can be determined from the reactivities of horizontal and vertical pools. Pooled peptides were diluted 10 times with water, followed by adding 1 μl of the solution into each corresponding well. S peptide pools and S_864-882_ peptide were also loaded as controls. S peptide pools of common cold HCoVs were loaded to assess cross-reactivities. For reporter cell assay, 100 μl of media containing T cells and APCs were added to the wells so that T cells are stimulated with each peptide at 1 μg/ml. After 20 hours, GFP reporter induction was assessed in T cell-gated population using AttuneNxT flow cytometer (Thermo Fisher Scientific).

### Restricting HLA determination

For the clonotypes detected in more than one donor from our sample pool, reporter cells were stimulated in the presence of HEK293T cells expressing individual pair of alleles that shared by the donors. Otherwise, reporter cells were stimulated in the presence of transformed B cells from various donors that have partial overlapped HLAs with the original donor of the clonotype.

### Tetramer staining

Freshly-isolated or *in vitro*-expanded PBMCs from DRB1*15:01 donors were stained with S_867-881_ peptide-loaded DRA-DRB1*15:01 tetramer. For expansion, PBMCs from DRB1*15:01-possessing donors were stimulated with 1 μg/ml of peptide S_864-882_ for 9 days. Cells were treated with 100 nM protease kinase inhibitor (dasatinib, Axon Medchem) at 37°C for 30 min and stained with tetramer conjugated with PE (Medical & Biological Laboratories) at 4°C for 1 hour. Cells were then stained with biotin-conjugated anti-PE antibody together with antibodies for surface markers for flow cytometric analysis using AttuneNxT (Thermo Fisher Scientific) and FlowJo 10.5.2 (BD Biosciences).

### Database analyses

Public repertoire datasets of pre-pandemic healthy donors (n = 786, Adaptive ImmuneACCESS)(Emerson et al., 2017) or unhospitalized COVID-19 convalescent donors who had mild symptoms (n = 1,413, Adaptive ImmuneRACE)(Nolan et al., 2020) of various ethnicities from US, Italy, and Spain were downloaded from the Adaptive Biotechnologies website and V gene names were renamed to match the IMGT nomenclature. Finally, clonotypes and their expansion were defined in the same way as in-house bulk TCR sequencing. TCR occurrence in a dataset was defined as the fraction of patients whose repertoire contained the given TCR, regardless of its expansion.

### Expression and purification of soluble TCRαβ heterodimer

Expression constructs encoding the extracellular domains of TCRα-017 (Q1-D207 in mature protein) and TCRβ-017 (G1-G241) subunits were incorporated into pCold vector including 6 × His-tag and a tobacco etch virus protease cleavage site. For crystallization, the point mutations (T159C in α and S166C/C184A in β) were introduced to form artificial disulfide bond as described previously (Boulter et al., 2003). The plasmids were transformed into *E. coli* BL21 competent cells Champion21 and Rosetta2 (DE3) (Novagen), respectively. The protein expression was induced by the addition of 0.5 mM IPTG at 18°C. Cells were suspended with 500 mM NaCl-containing Tris-HCl buffer (pH 8.0) and disrupted with sonication. The inclusion bodies including target proteins were collected by centrifugation. The inclusion body was then solubilized by 50 mM Tris-HCl buffer (pH 8.0) containing 6 M guanidine HCl, 10 mM EDTA and 2 mM DTT at room temperature. The equal amount of solubilized TCRα and TCRβ (35 mg each) was mixed and rapidly diluted with 1 L of 100 mM Tris-HCl buffer (pH 8.0) containing 5 M urea, 0.4 M L-arginine, 5 mM reduced glutathione and 0.5 mM oxidized glutathione at 4°C. The diluted solution was further dialyzed against 10 mM Tris-HCl buffer (pH 8.0) at 4°C for two days. The dialyzed solution was applied onto 5 ml Ni-NTA agarose (Fuji-film Wako) and His-tagged TCRαβ were eluted with elution buffer (50 mM Tris-HCl (pH 8.0), 300 mM NaCl and 250 mM imidazole). After removal of His-tag by TEV protease, the eluted protein was concentrated and further applied to Superdex75 (GE Healthcare) equilibrated with 20 mM Tris-HCl (pH 8.0) buffer containing 100 mM NaCl. The TCRαβ heterodimer fractions were concentrated up to 1.2 mg/ml by Amicon Ultra (molecular weight cut off: 10 kDa). The purity of the proteins was assessed by SDS-PAGE and CBB staining. The numbering of amino acids in CDR3 sequences of TCRs was based on mature protein.

### Biolayer interferometry analysis

Octet RED96 System (fortéBio, Pall Life Science) was used to detect the binding kinetics of TCRαβ heterodimer to peptide-loaded HLA molecule (Medical & Biological Laboratories). All assays were carried out at 25°C on a black 96-well plate with agitation set at 1,000 rpm and 200 μl/well for all solutions. Super Streptavidin biosensors (fortéBio, Pall Life Science) were dipped in PBS prior to use and reacted with 5.8 μg/ml biotinylated peptide-loaded HLA molecule. The association of TCRαβ heterodimer at concentrations of 0.1, 0.2 and 0.4 μM was measured for 900 sec, and the dissociation in PBS was measured for 900 sec. The obtained data was analyzed and processed using Octet data analysis 11.1 software.

### Crystallization, data collection, and structure determination of TCR-017 ectodomain

All crystallization trials were performed by sitting drop vapor diffusion method. Initial crystallization conditions were screened using Index (Hampton Research), SG1 Screen, and SG2 Screen (Molecular Dimensions). The best diffracted crystal was obtained under the condition of 0.1 M HEPES-Na (pH 7.0) and 1.1 M sodium malonate at 20°C. Prior to X-ray diffraction experiments, crystals were soaked in the reservoir containing 20% ethylene glycol and flash-cooled in liquid nitrogen. X-ray diffraction data sets were collected at the beamline BL-1A in the Photon Factory (Tsukuba, Japan). Diffraction data were integrated with program XDS (Kabsch, 2010) and scaled with program SCALA (Evans, 2006). The phases of datasets were determined by molecular replacement method using program MOLREP (Vagin and Teplyakov, 2010) with the coordinate of 1E6 TCR (PDB code: 5C0B)(Cole et al., 2016). After initial phase determination, the model buildings were manually performed using program COOT (Emsley et al., 2010). Refinement was performed using program REFMAC5 (Vagin et al., 2004) at the initial step and Phenix.refine for final model (Afonine et al., 2012). The stereochemical quality of the final model was assessed by program MolProbity (Williams et al., 2018). Data collection and refinement statistics were summarized in Table S7. Structural factors and the atomic coordinates of TCRαβ-017 ectodomain have been deposited in Protein Data Bank under the accession code of 7EA6. All figures of 3D structure were depicted with program PyMOL (The PyMOL Molecular Graphics System, Version 2.0 Schrödinger, LLC.).

### Antibodies

Anti-human CD3 (HIT3a), anti-human CD8 (RPA-T8), anti-human CD69 (FN50), anti-human CD137 (4B4-1), anti-human CD154 (24-31), anti-human CD4 (OKT4), anti-human CXCR5 (J252D4), TotalSeq-C Hashtags (LNH-94; 2M2), anti-mouse CD3 (17A2), anti-mouse CD69 (H1.2F3), anti-rat CD2 (OX-34) and biotin anti-phycoerythrin (PE)(PE001) antibodies were purchased from BioLegend. Rat IgG2b κ Isotype Control (eB149/10H5) was purchased from eBioscience.

### Statistical analysis

Statistical analysis was performed with *t*-test using SciPy and p-values were indicated in Figure 3A and 4D.

## Data availability

Recombinant SARS-CoV-2 S protein, TCR-reconstituted cells, soluble TCRαβ heterodimers under a standard material transfer agreement, and single cell-based transcriptome data are available from the corresponding author on request as supplies permit. The datasets supporting the current study have not been deposited in a public repository because permission was not acquired from the blood donors involved in this study but are available from the corresponding author on request. Other data needed to support the conclusion of this manuscript are included in the main text and Supplemental information.

## Supplemental Figure Legends

**Figure S1.**
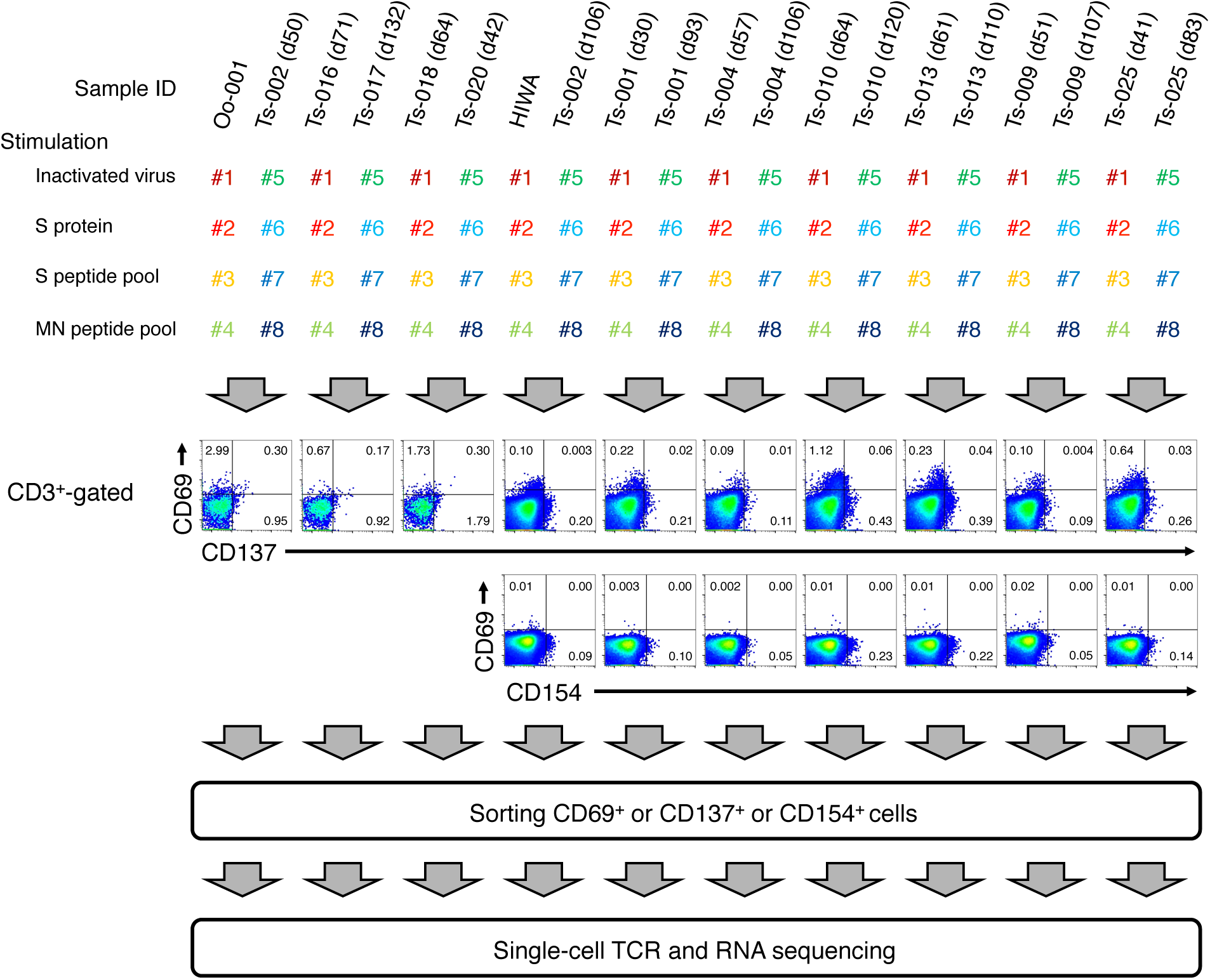
Workflow for single cell-based analyses of SARS-CoV-2-responsive T cells, related to Figure 1. PBMCs from healthy donors and convalescent COVID-19 patients were stimulated with SARS-CoV-2-derived antigens as indicated for 20 hours followed by staining with eight different hashtag antibodies (#1-8), anti-CD3 and indicated activation marker antibodies. Hashtagged cells from two patients were pooled and analyzed with cell sorter. Antigen-responded (CD69^+^ or CD137^+^ or CD154^+^) CD3^+^ T cells were sorted and immediately subjected to single-cell TCR and RNA sequencing. Each collection day after diagnosis is shown after the sample ID.

**Figure S2.**
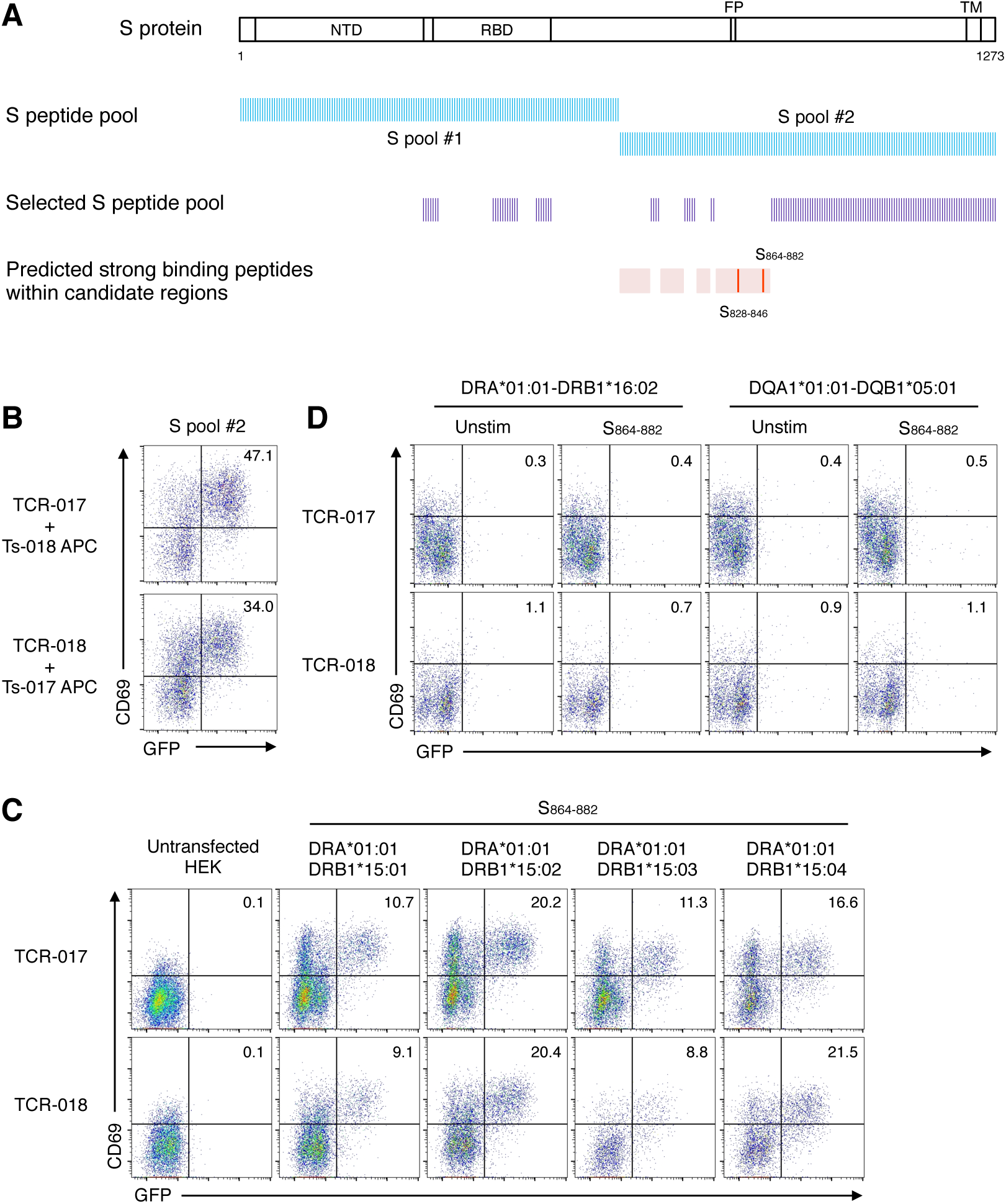
Epitope peptide and restricting HLAs of clonotype-017/018, related to Figure 2. (A) Relative positions that are covered by each S peptide pool are shown. NTD, N-terminal domain; RBD, receptor-binding domain; FP, fusion peptide; TM, transmembrane domain. Strong binding peptides (%RANK ≤ 2) presented by DRA-DRB1*15:01 in region 633-682, 708-740, 771-784 and 803-884 (pink regions) were predicted by NetMHC 4.0 Server. 19-mer peptides (red bars) that cover the strong binding peptides were synthesized and used for further epitope identification. (B) Cells expressing TCR-017 or -018 were stimulated with 1 μg/ml of S peptide pool #2 in the presence of heterologous APCs from exchanged donors (Ts-018 and Ts-017, respectively). (C and D) Cells expressing TCR-017 or -018 were stimulated with 1 μg/ml of peptide S_864-882_ in the presence of HEK293T cells expressing indicated HLAs (C) and immortalized B cells possessing indicated HLAs (D).

**Figure S3.**
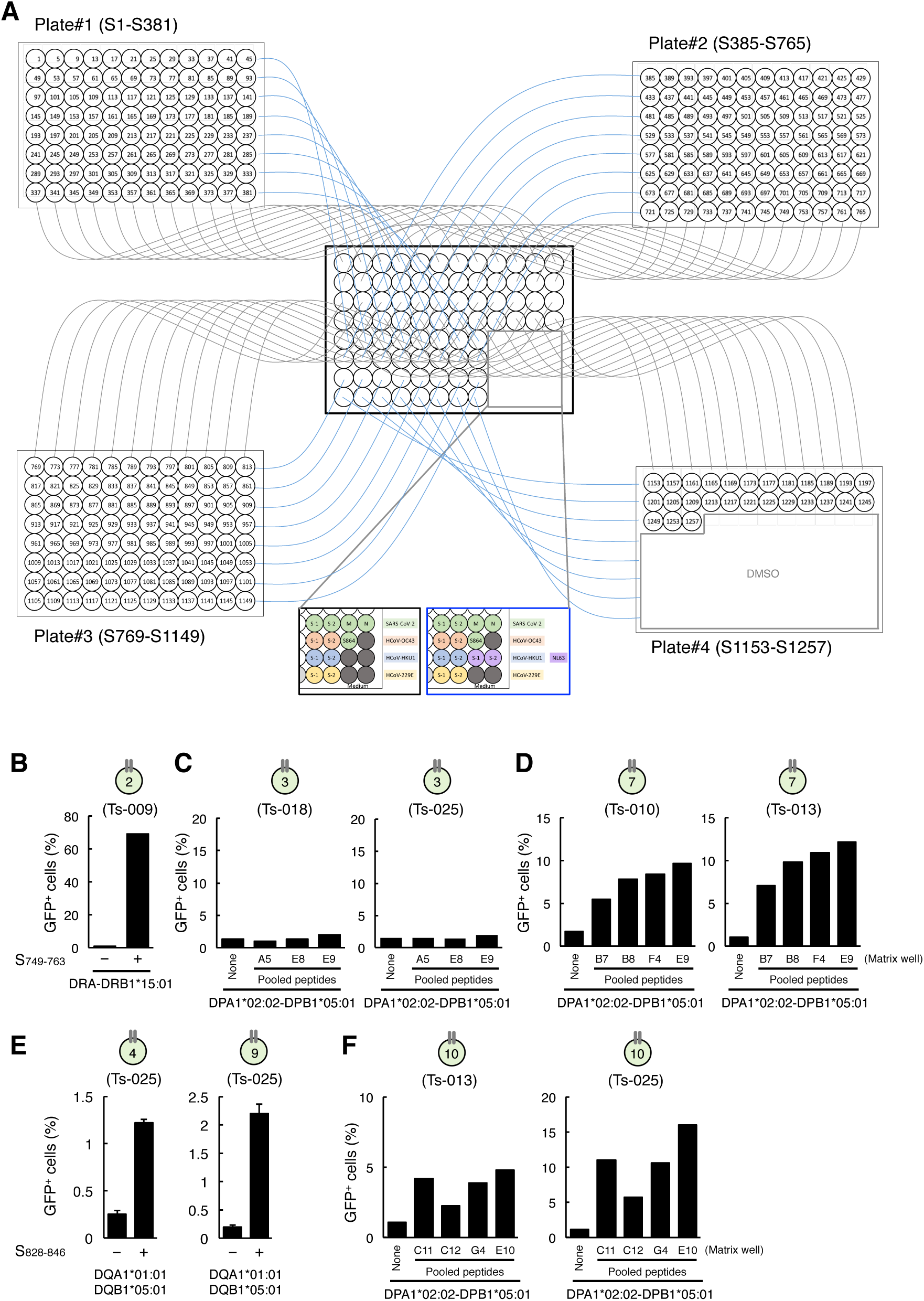
Identification of the epitopes of public clonotypes, related to Figure 3. (A) A scheme of rapid epitope-determination platform for identification of the epitope of T cell clonotypes. 15-mer peptides with 11 amino acids overlap that cover the full length of S protein of SARS-CoV-2 were synthesized and pooled as described in Materials and Methods. Each peptide is present in a unique combination of two different pools, thus the epitope can be revealed by the common peptide(s) in the pools that activated the clonotype. Peptide pools of S, M and N proteins of SARS-CoV-2 or HCoVs S proteins were also added in the matrix as controls. S864 indicates S_864-882_ peptide. (B) Identification of DRB1*15:01 as a restricting HLA of S_753-759_ for clonotype 2. Reporter cells expressing clonotype 2 were activated by S_749-763_ in the presence of HEK293T cells transfected with DRA*01:01-DRB1*15:01. Data are shown in mean ± SD of triplicates. (C) Exclusion of DPA1*02:02-DPB1*05:01 as a restricting HLA of S_353-367_ for clonotype 3. Reporter cells expressing clonotype 3 were not activated by S353-including peptide pools, such as A5, E8 and E9 (S pool #1) in Figure S3A, in the presence of HEK293T cells transfected with DPA1*02:02-DPB1*05:01. (D) Identification of DPA1*02:02-DPB1*05:01 as a restricting HLA of S_557-567_ for clonotype 7. Reporter cells expressing clonotype 7 were activated by S553- or S557-including peptide pools, such as B7, B8, F4 and E9 (S pool #1) in Figure S3A, in the presence of HEK293T cells transfected with DPA1*02:02-DPB1*05:01. (E) Identification of DQA1*01:01-DQB1*05:01 as a restricting HLA of S_833-843_ for clonotype 4 and 9. Reporter cells expressing clonotype 4 and 9 were activated by S_828-846_ in the presence of HEK293T cells transfected with DQA1*01:01-DQB1*05:01. Data are shown in mean ± SD of triplicates. (F) Identification of DPA1*02:02-DPB1*05:01 as a restricting HLA of S_957-967_ for clonotype 10. Reporter cells expressing clonotype 10 were activated by S953- or S957-including peptide pools, such as C11, C12, G4 and E10 (S pool #2) in Figure 3A, in the presence of HEK293T cells transfected with DPA1*02:02-DPB1*05:01.

**Figure S4.**
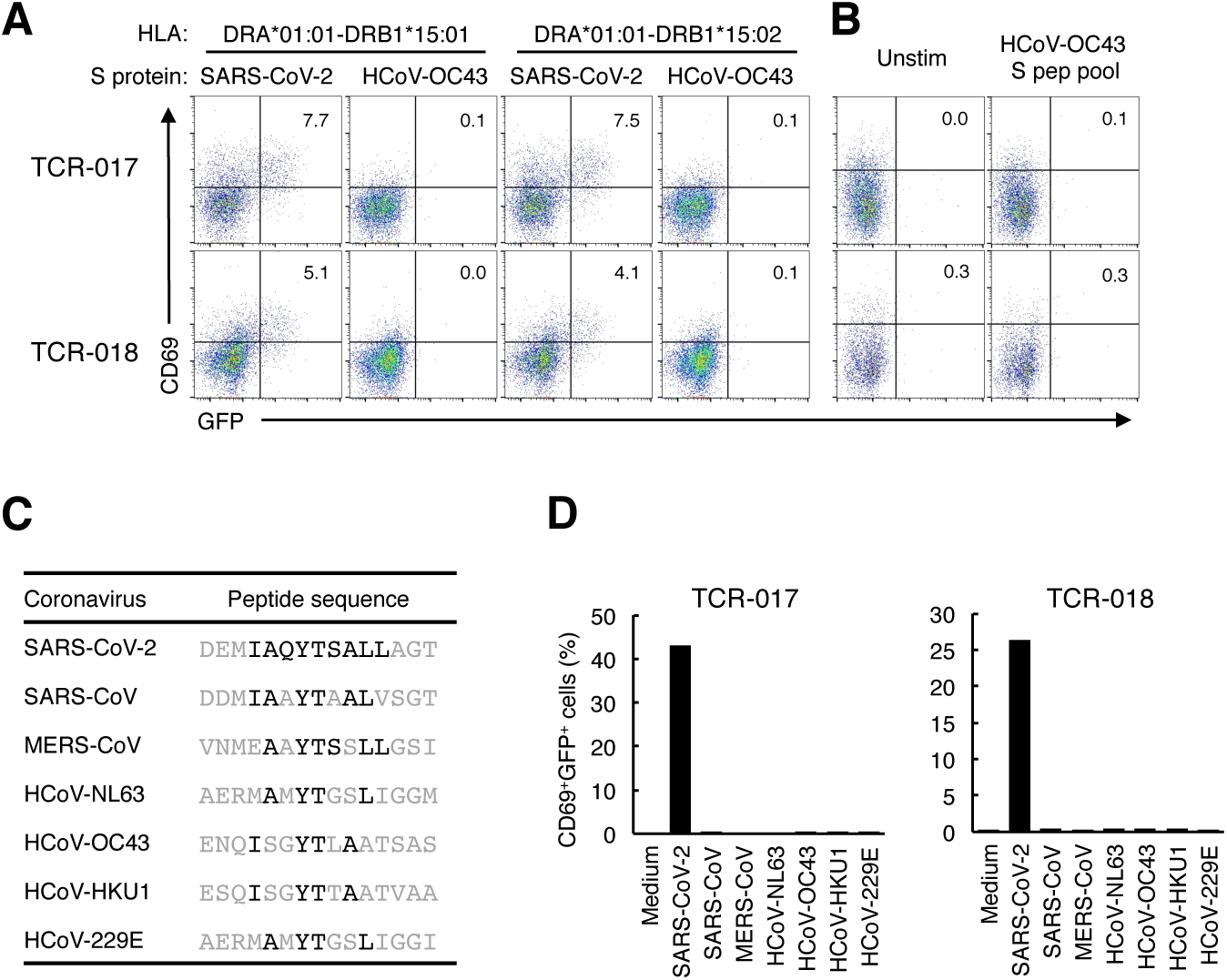
No cross-reactivity of clonotype-017/018 to other human coronaviruses, related to Figure 5. (A) Cells expressing TCR-017 or -018 were co-cultured with HEK293T cells expressing indicated HLAs and S protein (S) derived from indicated coronavirus. (B) Cells expressing TCR-017 or -018 were stimulated with 1 μg/ml of peptide pool derived from HCoV-OC43 S protein (HCoV-OC43 S pep pool). (C) Sequences of peptides from human coronaviruses corresponding to S_867-881_ of SARS-CoV-2. Amino acids that are same as those in SARS-CoV-2 in the core epitope are shown in black. (D) Cells expressing TCR-017 or -018 were stimulated with 1 μg/ml of peptides shown in (C).

**Figure S5.**
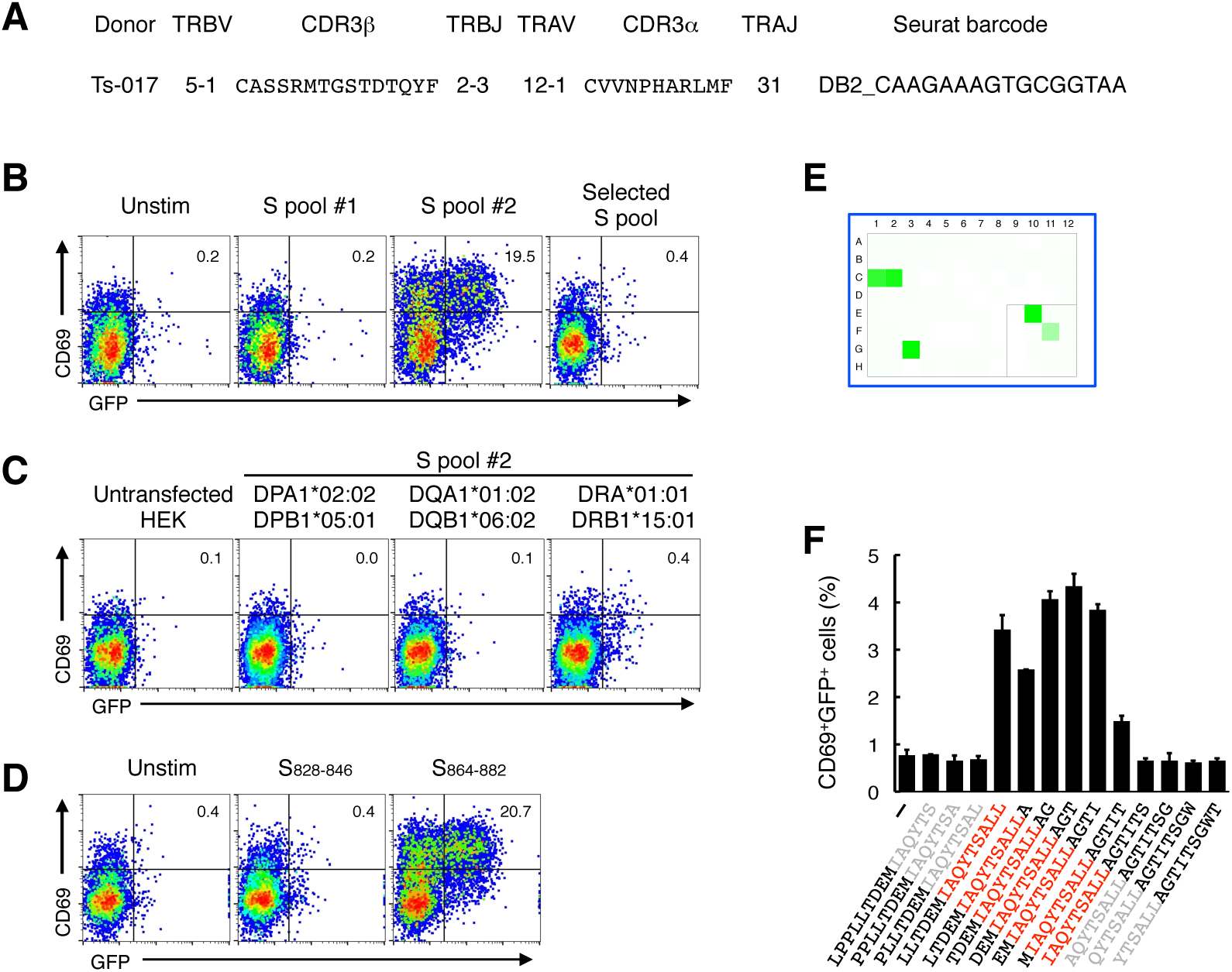
Another public Tfh clonotype that recognizes the same epitope S_870-878_ as clonotype-017/018, related to Figure 6. (A) V/J usage and CDR3 sequences of the newly-identified clonotype that recognize S_870-878_ peptide. (B-D) Reporter cells expressing TCRαβ in (A) was stimulated with 1 μg/ml of S peptide pool #1, #2 or selected S pool (S_304-338, 421-475, 492-519, 683-707, 741-770, 785-802, 885-1273_) (B), 1μg/ml of S peptide pool #2 in the presence of HEK293T cells expressing indicated HLAs (C), or 1 μg/ml of peptides S_828-846_ or S_864-882_ (D). (E) Cells were stimulated with pooled peptide matrix described in Figure S3A. Wells containing S865 or S869 (C1, C2 and G3), S pool #2 (E10) and S_864-882_ (F11) were positive. (F) 1 μg/ml of serial overlapped 15-mer peptides covering S_861-887_ region were tested for the reactivity to this clonotype to determine minimum epitope (shown in red). Data are shown in mean ± SD of triplicates.

**Figure S6.**
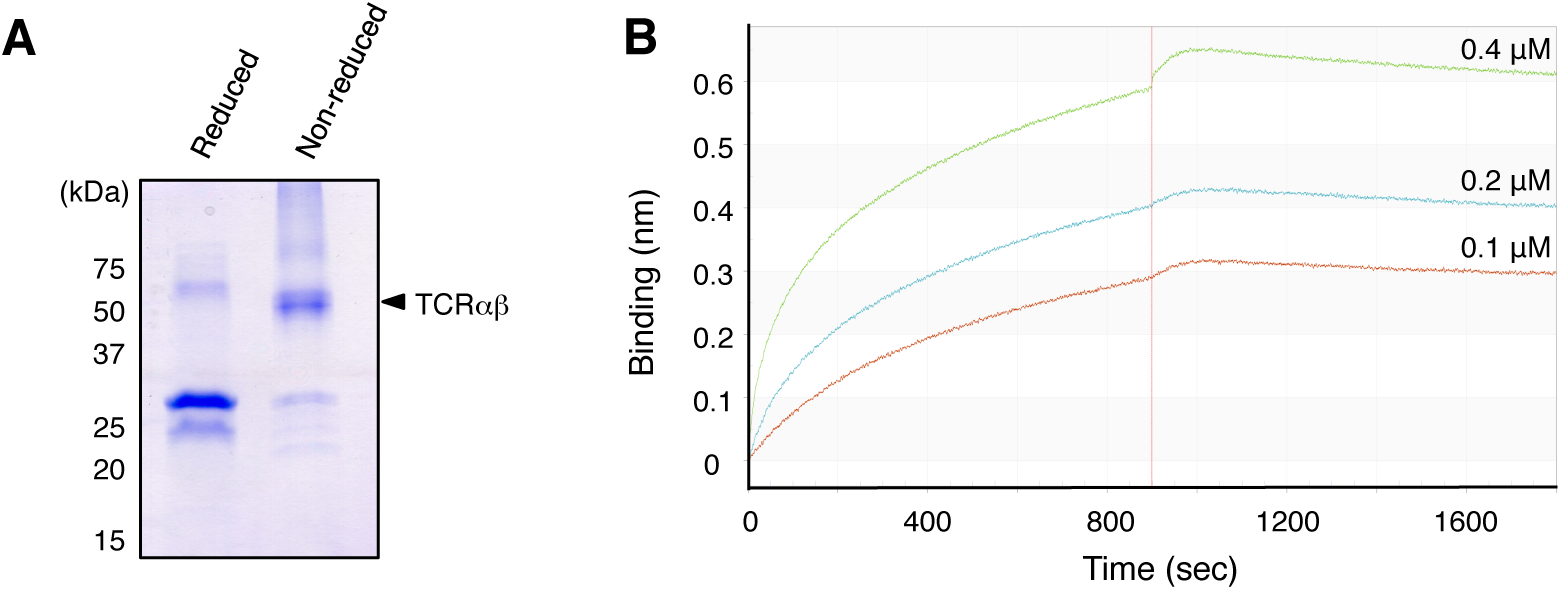
S_867-881_-loaded HLA-DR interacted with soluble TCR-017 heterodimer, related to Figure 6. (A) The purity of soluble TCR-017 αβ heterodimer (indicated by arrow head) was assessed by SDS-PAGE in reduced or non-reduced conditions. (B) Interaction between S_867-881_-loaded HLA DRA-DRB1*15:01 monomer and soluble TCR-017 by biolayer interferometry. Indicated concentrations of TCR αβ heterodimer were applied to 5.8 μg/ml immobilized recombinant HLA proteins.

**Table S1.**
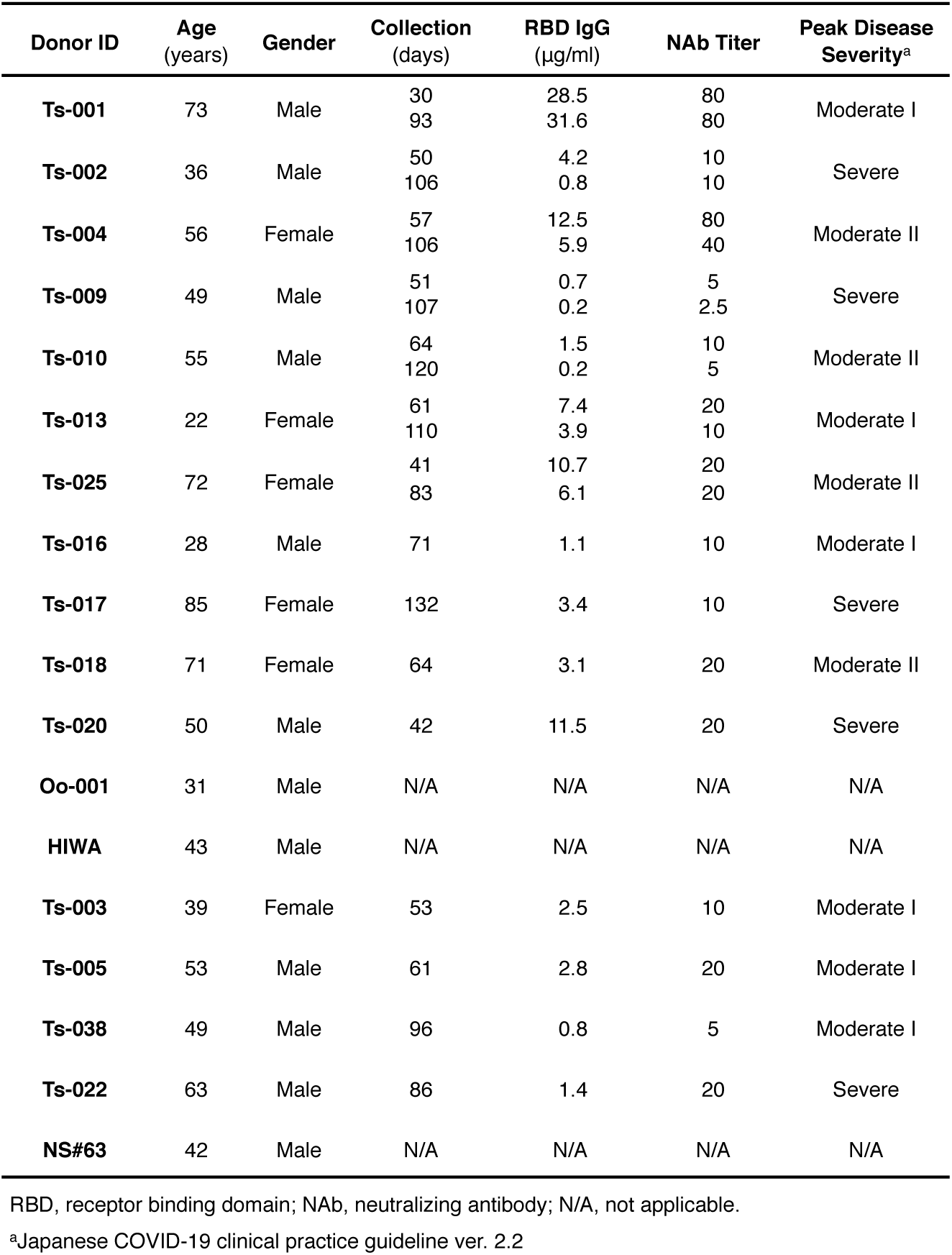
Donor characteristics, related to Figure 1.

| Donor ID | Age (years) | Gender | Collection (days) | RBD IgG (µg/ml) | NAb Titer | Peak Disease Severity<sup>a</sup> |
| --- | --- | --- | --- | --- | --- | --- |
| Ts-001 | 73 | Male | 30 | 28.5 | 80 | Moderate I |
|  |  |  | 93 | 31.6 | 80 |  |
| Ts-002 | 36 | Male | 50 | 4.2 | 10 | Severe |
|  |  |  | 106 | 0.8 | 10 |  |
| Ts-004 | 56 | Female | 57 | 12.5 | 80 | Moderate II |
|  |  |  | 106 | 5.9 | 40 |  |
| Ts-009 | 49 | Male | 51 | 0.7 | 5 | Severe |
|  |  |  | 107 | 0.2 | 2.5 |  |
| Ts-010 | 55 | Male | 64 | 1.5 | 10 | Moderate II |
|  |  |  | 120 | 0.2 | 5 |  |
| Ts-013 | 22 | Female | 61 | 7.4 | 20 | Moderate I |
|  |  |  | 110 | 3.9 | 10 |  |
| Ts-025 | 72 | Female | 41 | 10.7 | 20 | Moderate II |
|  |  |  | 83 | 6.1 | 20 |  |
| Ts-016 | 28 | Male | 71 | 1.1 | 10 | Moderate I |
| Ts-017 | 85 | Female | 132 | 3.4 | 10 | Severe |
| Ts-018 | 71 | Female | 64 | 3.1 | 20 | Moderate II |
| Ts-020 | 50 | Male | 42 | 11.5 | 20 | Severe |
| Oo-001 | 31 | Male | N/A | N/A | N/A | N/A |
| HIWA | 43 | Male | N/A | N/A | N/A | N/A |
| Ts-003 | 39 | Female | 53 | 2.5 | 10 | Moderate I |
| Ts-005 | 53 | Male | 61 | 2.8 | 20 | Moderate I |
| Ts-038 | 49 | Male | 96 | 0.8 | 5 | Moderate I |
| Ts-022 | 63 | Male | 86 | 1.4 | 20 | Severe |
| NS#63 | 42 | Male | N/A | N/A | N/A | N/A |

RBD, receptor binding domain; NAb, neutralizing antibody; N/A, not applicable.
<sup>a</sup>Japanese COVID-19 clinical practice guideline ver. 2.2

**Table S2.** HLA class II types of involved blood donors, related to Figure 1.

| Donor ID | DPA1 | DPB1 | DQA1 | DQB1 | DRB1 | DRB3/4/5 |
| --- | --- | --- | --- | --- | --- | --- |
| **Ts-001** | 02:02/02:02 | 05:01/05:01 | 03:01/03:02 | 03:02<sup>b</sup>/03:03 | 04:03/12:01<sup>c</sup> | 4*01:04/3*01:01 |
| **Ts-002** | 01:03/02:02 | 02:01/03:01 | 05:03/05:08 | 03:01<sup>d</sup>/03:01<sup>d</sup> | 12:01<sup>c</sup>/14:03 | 3*01:01/3*01:01 |
| **Ts-004** | 01:03/02:02 | 02:01/05:01 | 01:02/03:02 | 03:03/06:04 | 09:01/13:02 | 4*01:03/3*03:01 |
| **Ts-009** | 02:02/02:02 | 02:01/02:02 | 01:02/03:03 | 04:01/06:02 | 04:05/15:01 | 4*01:03/5*01:01 |
| **Ts-010** | 02:01/02:02 | 05:01/09:01 | 01:03/01:03 | 06:01/06:01 | 08:03/15:02<sup>e</sup> | –<sup>a</sup>/5*01:02 |
| **Ts-013** | 01:03/02:02 | 04:02/05:01 | 01:02/03:03 | 04:02/05:02<sup>f</sup> | 04:04/16:02 | 4*01:03/5*02:02 |
| **Ts-025** | 01:03/02:02 | 04:02/05:01 | 01:01/03:01 | 03:02<sup>b</sup>/05:01 | 01:01/04:03 | –<sup>a</sup>/4*01:03 |
| **Ts-016** | 01:03/02:02 | 02:01/05:01 | 01:03/05:08 | 03:01<sup>d</sup>/06:01 | 12:01<sup>c</sup>/15:02<sup>e</sup> | 3*01:01/5*01:02 |
| **Ts-017** | 02:02/02:02 | 02:01/05:01 | 01:02/01:02 | 06:02/06:04 | 13:02/15:01 | 3*03:01/5*01:01 |
| **Ts-018** | 02:02/02:02 | 03:01/05:01 | 01:02/03:02 | 03:03/06:02 | 09:01/15:01 | 4*01:03/5*01:01 |
| **Ts-020** | 02:02/02:02 | 05:01/05:01 | 01:02/01:03 | 06:01/06:02 | 08:03/15:01 | 5*01:01/–<sup>a</sup> |
| **Oo-001** | 01:03/02:02 | 02:01/04:02 | 01:01/03:01 | 03:02<sup>b</sup>/05:01 | 01:01/04:03 | 4*01:03/–<sup>a</sup> |
| **HIWA** | 01:03/02:02 | 02:01/05:01 | 01:05/03:03 | 03:01<sup>d</sup>/05:01 | 04:01/10:01 | 4*01:02/–<sup>a</sup> |
| **Ts-003** | 02:02/02:02 | 05:01/05:01 | 01:02/05:05 | 03:01<sup>g</sup>/05:02<sup>f</sup> | 11:01/15:02<sup>e</sup> | 3*02:02/5*01:01 |
| **Ts-005** | 02:02/02:02 | 02:01/05:01 | 01:02/05:07 | 03:01<sup>d</sup>/06:02 | 14:03/15:01 | 3*01:01/5*01:01 |
| **Ts-038** | 01:03/02:01 | 02:01<sup>h</sup>/09:01<sup>i</sup> | 01:02/01:03 | 06:01/06:04 | 13:02/15:02<sup>e</sup> | 3*03:01/5*01:02 |
| **Ts-022** | 01:03/02:02 | 02:01/05:01 | 01:03/01:04 | 05:03/06:01 | 14:54/15:02<sup>e</sup> | 3*02:02/5*01:02 |
| **NS#63** | 02:01/02:01 | 09:01/09:01 | 01:02/01:03 | 06:01/06:02 | 15:01/15:02<sup>e</sup> | 5*01:01/5*01:02 |

<sup>a</sup>A dash (–) indicates no allele expressed.
<sup>b–i</sup>We cannot exclude <sup>b</sup>DQB1 03:289/03:416, <sup>c</sup>DRB1 12:10, <sup>d</sup>DQB1 03:276N, <sup>e</sup>DRB1 15:140/15:149,
<sup>f</sup>DQB1 05:241, <sup>g</sup>DQB1 03:297/03:419, <sup>h</sup>DPB1 352:01 and <sup>i</sup>DPB1 899:01.

**Table S3. Tfh clonotypes identified in single cell analysis, related to Figure 1-3.**

**Table S4.** Proteins possessing epitopes of TCR-017/018 (IAQYTSAAL), related to Figure 5.

| Cross-reactive protein [Organism] | Identity (%) | Accession |
| --- | --- | --- |
| histidine ammonia-lyase [*Haloarcula salaria*] | 100 | WP_188853020.1 |
| histidine ammonia-lyase [*Haloarcula argentinensis*] | 100 | WP_005537844.1 |
| histidine ammonia-lyase [*Halorubellus* sp. JP-L1] | 100 | WP_166041822.1 |
| histidine ammonia-lyase [*Deinococcus wulumuqiensis*] | 100 | WP_114672982.1 |
| histidine ammonia-lyase [*Deinococcus fonticola*] | 100 | WP_135229009.1 |
| histidine ammonia-lyase [*Deinococcus radiodurans*] | 100 | WP_010889407.1 |
| histidine ammonia-lyase [*Deinococcus wulumuqiensis*] | 100 | WP_017871478.1 |
| histidine ammonia-lyase [*Deinococcus* sp. K2S05-167] | 100 | WP_119766766.1 |
| TPA: histidine ammonia-lyase [*Ignavibacteriales* bacterium] | 100 | HCY74963.1 |
| histidine ammonia-lyase [*Leptolyngbya valderiana*] | 100 | WP_063716510.1 |
| histidine ammonia-lyase [*Anaerolineae* bacterium] | 100 | NPV66375.1 |
| histidine ammonia-lyase [*Candidatus* Parcubacteria bacterium] | 100 | PLX26207.1 |
| histidine ammonia-lyase [*Candidatus* Parcubacteria bacterium] | 100 | RJQ35361.1 |
| Histidine ammonia-lyase [*Synergistetes* bacterium ADurb.BinA166] | 100 | OPZ34295.1 |
| histidine ammonia-lyase [*Candidatus* Fermentibacter daniensis] | 100 | NLI03160.1 |
| hypothetical protein AO395_09060 [*Candidatus* Fermentibacter daniensis] | 100 | KZD18851.1 |
| histidine ammonia-lyase [*Aminivibrio pyruvatiphilus*] | 100 | TDY64917.1 |
| aromatic amino acid lyase [*Sciscionella marina*] | 100 | WP_020501521.1 |
| histidine ammonia-lyase [*Synergistaceae* bacterium] | 100 | NLM72193.1 |
| aromatic amino acid lyase [*Sciscionella* sp. SE31] | 100 | WP_031466349.1 |
| histidine ammonia-lyase [*Pseudomonas* sp. CMR12a] | 100 | WP_124346689.1 |
| histidine ammonia-lyase [*Gammaproteobacteria* bacterium] | 100 | PCI68862.1 |
| histidine ammonia-lyase [*Kangiella* sp.] | 100 | PHS17023.1 |
| histidine ammonia-lyase [*Candidatus* Lokiarchaeota archaeon] | 100 | NHJ25594.1 |
| histidine ammonia-lyase [*Candidatus* Lokiarchaeota archaeon] | 100 | MBD3194455.1 |
| histidine ammonia-lyase [*Deinococcus proteolyticus*] | 100 | WP_013622983.1 |
| histidine ammonia-lyase [*Deinococcus piscis*] | 100 | WP_189641912.1 |
| histidine ammonia-lyase [*Synergistales* bacterium] | 100 | NCC56342.1 |
| histidine ammonia-lyase [*Aminivibrio pyruvatiphilus*] | 100 | WP_133955739.1 |
| hypothetical protein AO394_00950 [*Candidatus* Fermentibacter daniensis] | 100 | KZD16260.1 |
| histidine ammonia-lyase [*Synergistaceae* bacterium] | 100 | NLD96450.1 |
| histidine ammonia-lyase [*Synergistales* bacterium] | 100 | NCB17450.1 |
| histidine ammonia-lyase [*Deltaproteobacteria* bacterium HGW-Deltaproteobacteria-20] | 100 | PKN47942.1 |
| Histidine ammonia-lyase [candidate division Hyd24-12 bacterium ADurb.Bin004] | 100 | OQC66456.1 |
| aromatic amino acid lyase [*Synergistaceae* bacterium] | 100 | NLK18261.1 |
| aromatic amino acid lyase [*Phycisphaerales* bacterium] | 100 | NRA57503.1 |
| Histidine ammonia-lyase [*Synergistetes* bacterium ADurb.BinA166] | 100 | OPZ38667.1 |

**Table S5.**
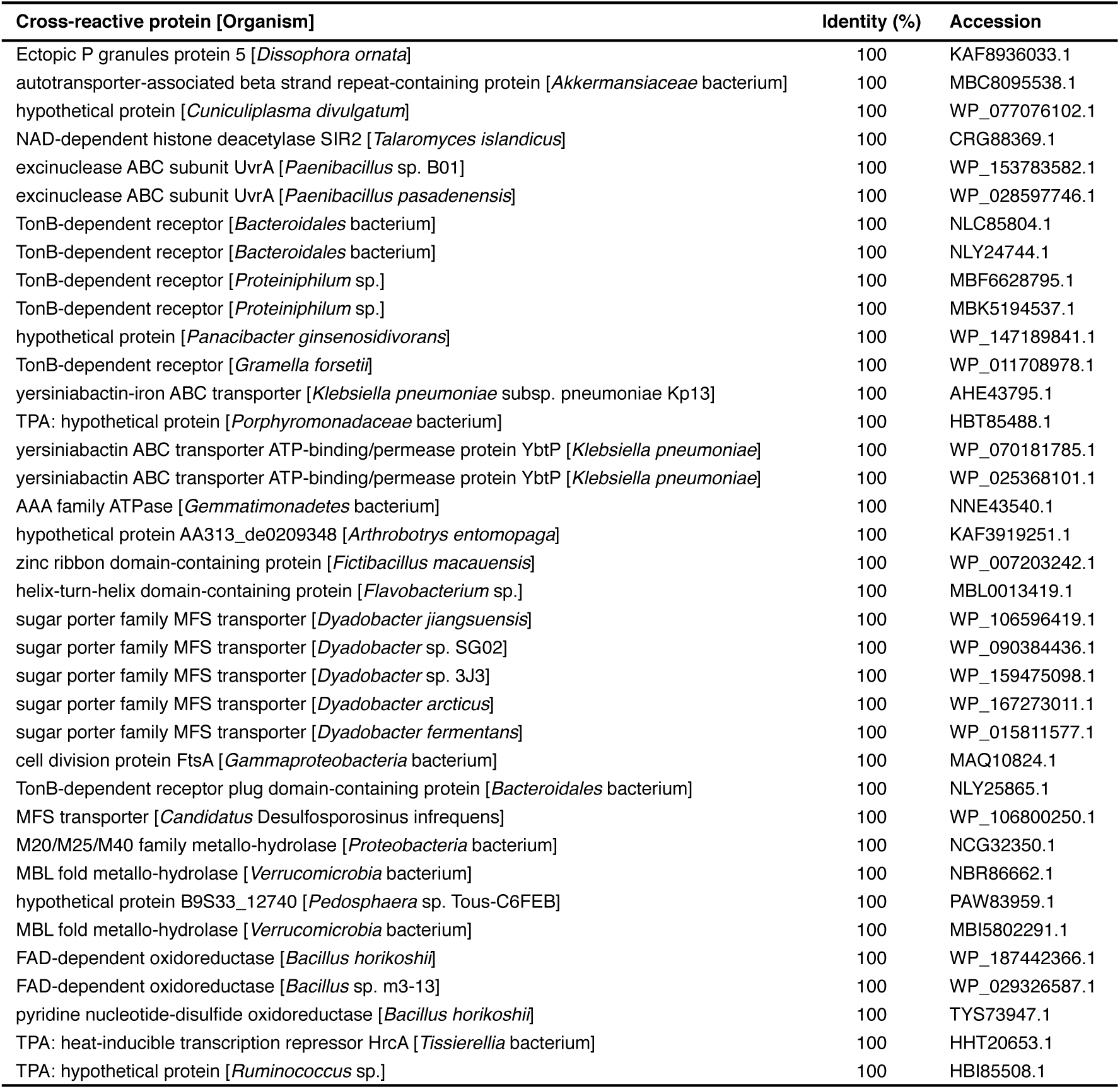
Proteins possessing epitopes of clonotype 2 (LLQYGSF), related to Figure 5.

| Cross-reactive protein [Organism] | Identity (%) | Accession |
| --- | --- | --- |
| Ectopic P granules protein 5 [*Dissophora ornata*] | 100 | KAF8936033.1 |
| autotransporter-associated beta strand repeat-containing protein [*Akkermansiaceae* bacterium] | 100 | MBC8095538.1 |
| hypothetical protein [*Cuniculiplasma divulgatum*] | 100 | WP_077076102.1 |
| NAD-dependent histone deacetylase SIR2 [*Talaromyces islandicus*] | 100 | CRG88369.1 |
| excinuclease ABC subunit UvrA [*Paenibacillus* sp. B01] | 100 | WP_153783582.1 |
| excinuclease ABC subunit UvrA [*Paenibacillus pasadenensis*] | 100 | WP_028597746.1 |
| TonB-dependent receptor [*Bacteroidales* bacterium] | 100 | NLC85804.1 |
| TonB-dependent receptor [*Bacteroidales* bacterium] | 100 | NLY24744.1 |
| TonB-dependent receptor [*Proteiniphilum* sp.] | 100 | MBF6628795.1 |
| TonB-dependent receptor [*Proteiniphilum* sp.] | 100 | MBK5194537.1 |
| hypothetical protein [*Panacibacter ginsenosidivorans*] | 100 | WP_147189841.1 |
| TonB-dependent receptor [*Gramella forsetii*] | 100 | WP_011708978.1 |
| yersiniabactin-iron ABC transporter [*Klebsiella pneumoniae* subsp. *pneumoniae* Kp13] | 100 | AHE43795.1 |
| TPA: hypothetical protein [*Porphyromonadaceae* bacterium] | 100 | HBT85488.1 |
| yersiniabactin ABC transporter ATP-binding/permease protein YbtP [*Klebsiella pneumoniae*] | 100 | WP_070181785.1 |
| yersiniabactin ABC transporter ATP-binding/permease protein YbtP [*Klebsiella pneumoniae*] | 100 | WP_025368101.1 |
| AAA family ATPase [*Gemmatimonadetes* bacterium] | 100 | NNE43540.1 |
| hypothetical protein AA313_de0209348 [*Arthrobotrys entomopaga*] | 100 | KAF3919251.1 |
| zinc ribbon domain-containing protein [*Fictibacillus macauensis*] | 100 | WP_007203242.1 |
| helix-turn-helix domain-containing protein [*Flavobacterium* sp.] | 100 | MBL0013419.1 |
| sugar porter family MFS transporter [*Dyadobacter jiangsuensis*] | 100 | WP_106596419.1 |
| sugar porter family MFS transporter [*Dyadobacter* sp. SG02] | 100 | WP_090384436.1 |
| sugar porter family MFS transporter [*Dyadobacter* sp. 3J3] | 100 | WP_159475098.1 |
| sugar porter family MFS transporter [*Dyadobacter arcticus*] | 100 | WP_167273011.1 |
| sugar porter family MFS transporter [*Dyadobacter fermentans*] | 100 | WP_015811577.1 |
| cell division protein FtsA [*Gammaproteobacteria* bacterium] | 100 | MAQ10824.1 |
| TonB-dependent receptor plug domain-containing protein [*Bacteroidales* bacterium] | 100 | NLY25865.1 |
| MFS transporter [*Candidatus Desulfosporosinus infrequens*] | 100 | WP_106800250.1 |
| M20/M25/M40 family metallo-hydrolase [*Proteobacteria* bacterium] | 100 | NCG32350.1 |
| MBL fold metallo-hydrolase [*Verrucomicrobia* bacterium] | 100 | NBR86662.1 |
| hypothetical protein B9S33_12740 [*Pedosphaera* sp. Tous-C6FEB] | 100 | PAW83959.1 |
| MBL fold metallo-hydrolase [*Verrucomicrobia* bacterium] | 100 | MBI5802291.1 |
| FAD-dependent oxidoreductase [*Bacillus horikoshii*] | 100 | WP_187442366.1 |
| FAD-dependent oxidoreductase [*Bacillus* sp. m3-13] | 100 | WP_029326587.1 |
| pyridine nucleotide-disulfide oxidoreductase [*Bacillus horikoshii*] | 100 | TYS73947.1 |
| TPA: heat-inducible transcription repressor HrcA [*Tissierellia* bacterium] | 100 | HHT20653.1 |
| TPA: hypothetical protein [*Ruminococcus* sp.] | 100 | HBI85508.1 |

**Table S6.**
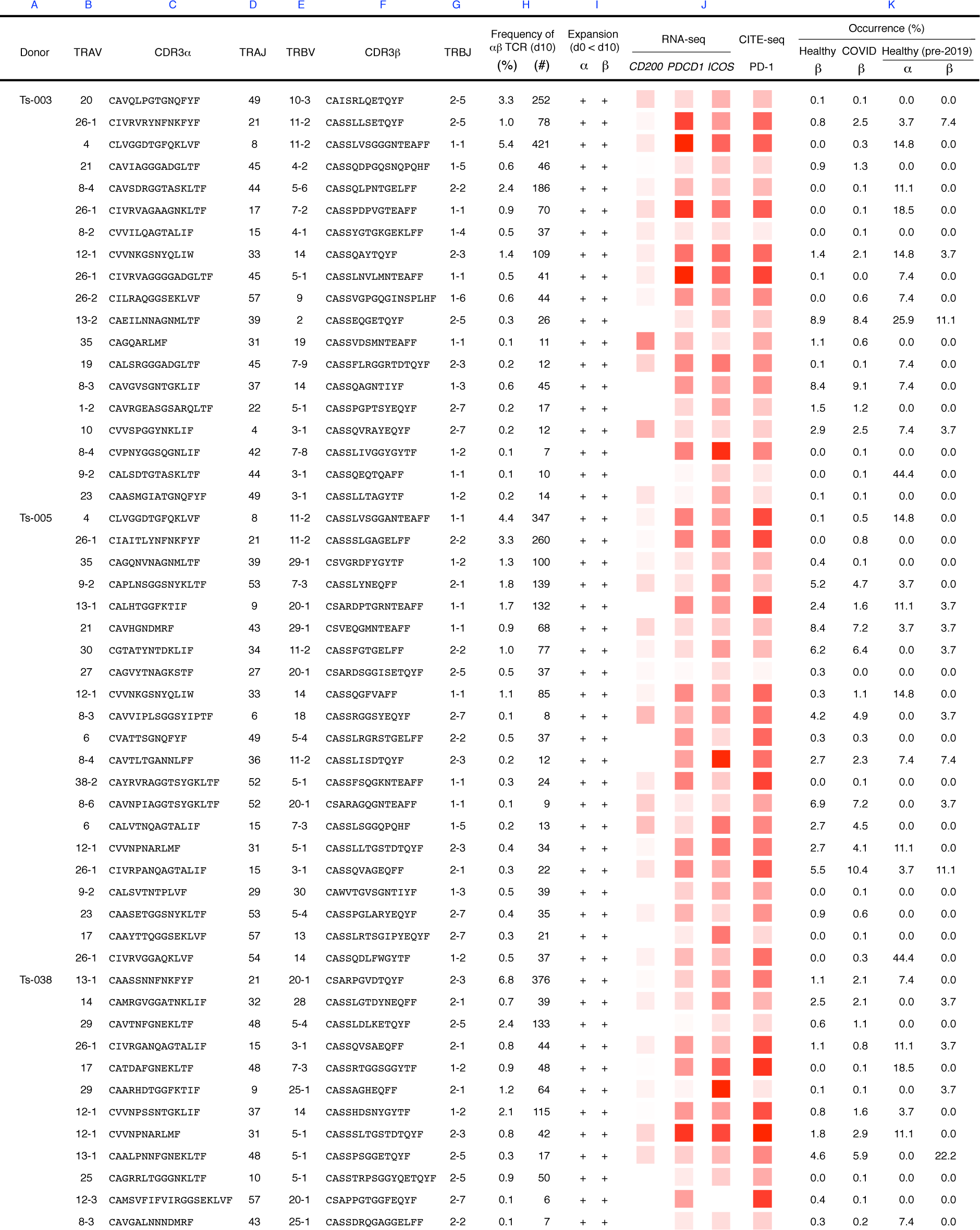
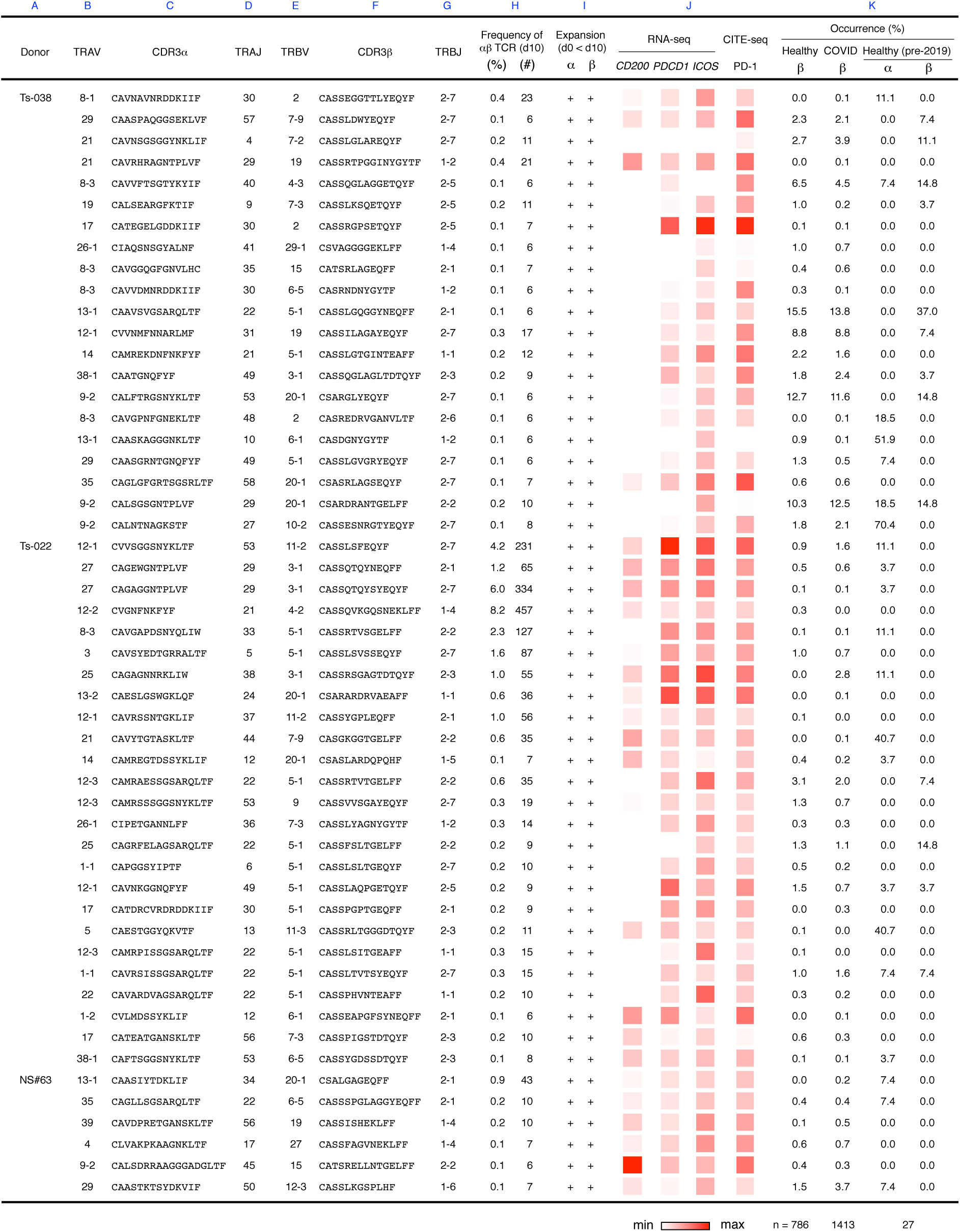
S_864-882_-reacted public Tfh clonotypes, related to Figure 6. Clonotypes excluding those with Treg characteristics were assembled with in-house bulk TCR-seq data as well as those in cohort databases. Clonotypes that satisfied following two criteria are shown: 1) expanded during stimulation based on single cell sequencing (> 0.1% of total CD4^+^ T cells) and 2) detected in cohort databases.

**Table S7.**
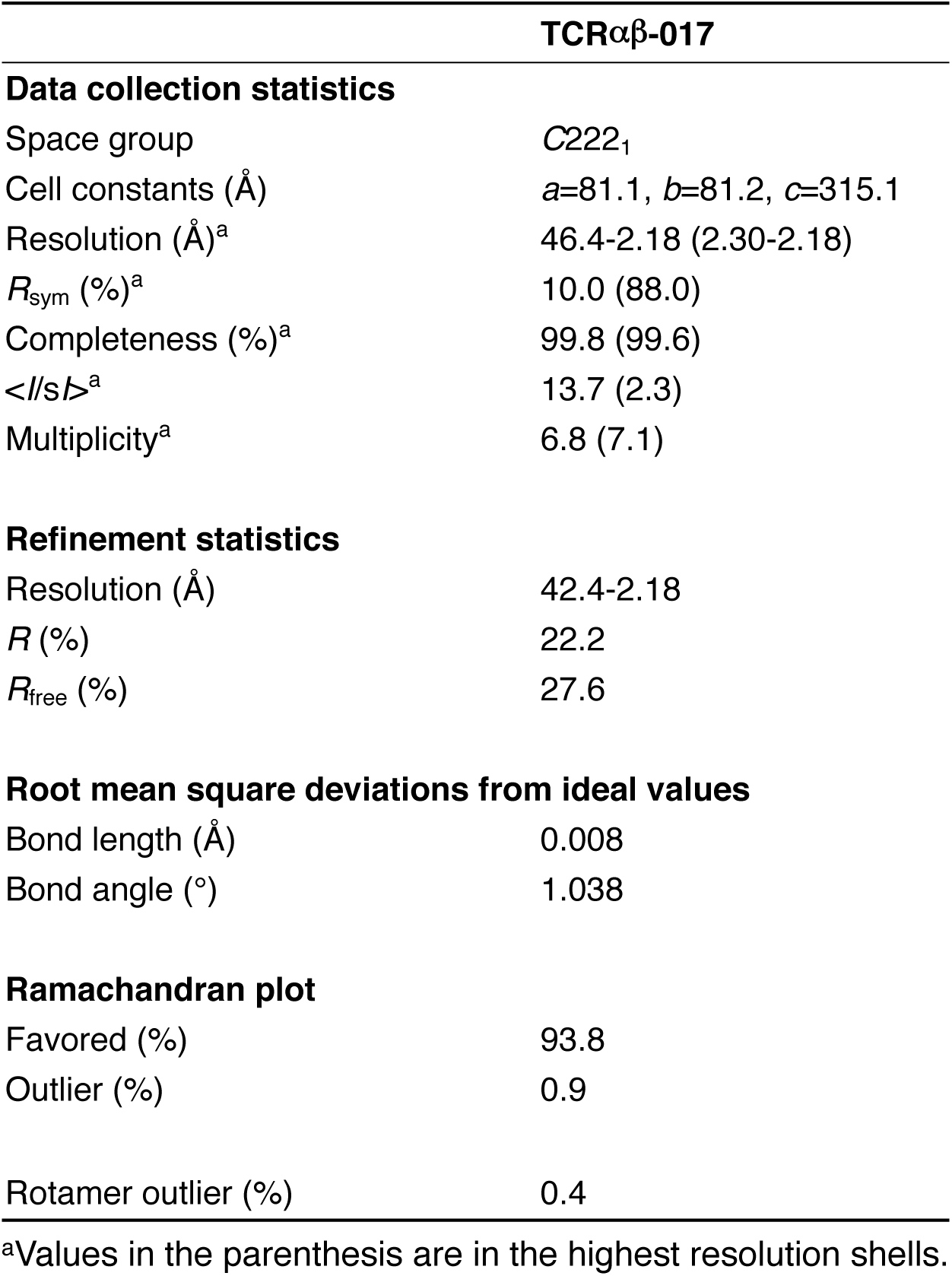
Data collection and refinement statistics of the crystallographic analysis of TCRαβ-017, related to Figure 4.

